# Western diet unmasks transient low-level vinyl chloride exposure-induced tumorigenesis; potential role of the (epi-)transcriptome

**DOI:** 10.1101/2022.02.24.481808

**Authors:** Silvia Liu, Liqing He, Olivia B Bannister, Regina D Schnegelberger, Junyan Tao, Andrew D Althouse, Francisco J Schopfer, Banrida Wahlang, Matthew C Cave, Satdarshan P Monga, Xiang Zhang, Gavin E Arteel, Juliane I Beier

**Affiliations:** Department of Pathology, University of Pittsburgh; Pittsburgh Liver Research Center, Pittsburgh, PA 15213; Department of Chemistry, University of Louisville, Louisville, KY 40208; Department of Medicine, Division of Gastroenterology, Hepatology and Nutrition; University of Pittsburgh; Department of Pharmacology and Chemical Biology, University of Pittsburgh; Division of General Internal Medicine, University of Pittsburgh, Pittsburgh, PA 15213; Division of Gastroenterology, Hepatology and Nutrition, Department of Medicine, University of Louisville School of Medicine, Louisville, KY, 40202, USA; Superfund Research Center, University of Louisville, Louisville, KY, 40202, USA; Hepatobiology and Toxicology Center, University of Louisville, Louisville, KY, 40202, USA; Center for Integrative Environmental Health Sciences, University of Louisville, Louisville, KY, 40202, USA; University of Louisville Alcohol Research Center, Louisville, KY, 40202, USA; Department of Pharmacology and Toxicology, University of Louisville School of Medicine, Louisville, KY, 40202; Department of Biochemistry and Molecular Genetics, University of Louisville School of Medicine, Louisville, KY 40202; Liver Transplant Program at UofL Health-Jewish Hospital Trager Transplant Center, Louisville, KY, 40202, USA; The Robley Rex Veterans Affairs Medical Center, Louisville, KY, 40206, USA; Department of Environmental and Occupational Health; University of Pittsburgh, Pittsburgh, PA 15213

**Keywords:** chloroethylene, volatile organic compound, high-fat diet, exposome, transcriptome, epitranscriptome, hepatocellular cancer

## Abstract

**Background & Aims:** Vinyl chloride (VC) monomer is a volatile organic compound commonly used in industry to produce polyvinyl chloride (PVC). At high exposure levels, VC causes liver cancer and toxicant-associated steatohepatitis. However, lower exposure levels (i.e., < regulatory exposure limits) that do not directly damage the liver, enhance injury caused by Western diet (WD). Although these lower exposure levels are considered ‘safe,’ it is unknown if the long-term impact of transient low-concentration VC enhances the risk of liver cancer development. Low-concentration VC is especially a concern given that fatty liver disease is in and of itself a risk factor for the development of liver cancer.

**Methods:** C57Bl/6J mice were fed WD or control diet (CD) for 1 year. During the first 12 weeks of feeding only, mice were also exposed to VC via inhalation at sub-regulatory limit concentrations (<1 ppm) or air for 6 hours/day, 5 days/week.

**Results:** Feeding WD for 1 year caused significant hepatic injury, including steatohepatitis and moderate fibrosis, which was exacerbated by VC. Additionally, VC increased the number of tumors which ranged from moderately to poorly differentiated hepatocellular carcinoma (HCC). Transcriptomic analysis demonstrated VC-induced changes in metabolic but also ribosomal processes. Epitranscriptomic analysis showed a VC-induced shift of the modification pattern that has been associated with metabolic disease, mitochondrial dysfunction, and cancer.

**Conclusions:** These data indicate that VC sensitizes the liver to other stressors (e.g., WD), resulting in enhanced tumorigenesis. These data raise concerns about a potential interaction between VC exposure and WD. Furthermore, it also emphasizes that current safety restrictions may be insufficient to account for other factors that can influence hepatotoxicity.

## Introduction

The global burden of chronic liver disease (CLD) has been steadily increasing.[1, 2] CLD is not a single pathological manifestation but rather a spectrum with various stages of severity, ranging from steatosis to steatohepatitis and subsequently to the more severe fibrosis and cirrhosis.[3] The incidence of HCC has also been increasing in parallel with that of CLD,[4, 5] as ~90% of HCC occurs on the background of advanced CLD, especially cirrhosis.[6, 7] HCC is the 6^th^ most common cancer worldwide.[4] However, as treatment options for advanced HCC are limited, the 5-year survival rate for HCC is low, and it is consequently the 2^nd^ most lethal solid cancer.[8] Unlike most cancers, the mortality rate of HCC has only improved incrementally in recent decades.[9]

Liver diseases are well-known to show categorical health disparities within a given population.[10–12] One of the components of consideration in local health disparities is differential exposure to environmental chemicals.[13] Indeed, disparities in exposure to environmental hazards often overlap with other sociodemographic inequities.[14] Although severe CLD is the primary risk factor for the development of HCC, developing studies indicate that overall risk is modified by environmental factors. The prevailing hypothesis is that these risk modifiers act by increasing the mutation frequency in the damaged liver and/or by favoring clonal expansion and progression of precancerous lesions. Moreover, exposure to a myriad of anthropogenic chemicals has been linked with an increase in HCC incidence, including vinyl chloride (VC).[15–18]

VC is a ubiquitous environmental pollutant. Its global production was recently estimated at 27 million metric tons annually.[19] VC has been identified as a solvent degradation product at waste sites, in the groundwater near military installations and natural gas fracking sites.[20, 21] Major environmental exposure risk stems from ambient air (concentrations of VC near production sites can be quite high - PPM range), or from contaminated groundwater.[22] VC readily volatilizes in buildings located above contaminated groundwater, where these vapors then accumulate.[23, 24] In support of the ubiquity of VC exposure, a recent study indicated that neonates have already adult exposure levels to VC and other VOCs.[25] Owing to its widespread presence and its known potential human risk, *VC is ranked #4 on the ATSDR Hazardous Substance Priority List*.[26]

Exposure to some anthropogenic chemicals is associated with increased HCC risk, including VC.[27] However, their prevalence is inadequately quantified, and their epidemiology limited. Moreover, whether this is a direct effect on HCC development or an indirect effect (via causing CLD) is not clear and the diagnosis of liver injury caused by chemicals is challenging, one of exclusion and often requires an interdisciplinary approach.[28] Most research has focused on high occupational exposure to VC (>1 ppm). [29–31] However, recent data demonstrate that VC, at concentrations that are not hepatotoxic per se, exacerbates experimental NAFLD.[32, 33] Moreover, VC is an independent risk factor for liver cancer in patients with CLD from other causes (e.g., alcohol, HCV, etc), suggesting that VC exposure may enhance HCC risk.[34] Therefore, the major goal of the study was to prove this hypothesis using this newly developed animal model of exposure to low levels of VC in combination with a Western diet.

## Materials and methods

### Animals and Treatments

Six-week-old male C57Bl/6J mice were purchased from Jackson Laboratory (Bar Harbor, ME). Mice were housed in a pathogen-free barrier accredited by the Association for Assessment and Accreditation of Laboratory Animal Care, and procedures were approved by the local Institutional Animal Care and Use Committee. Food and tap water were provided ad libitum. Mice were fed either control diet (CD, 13% fat, Teklad diets, #TD.120336) or Western-style diet (WD, 42% fat, Teklad diets, #TD.07511) for 12 weeks. Mice were exposed to VC (Kin-tek, La Marque, TX) at ~0.85 ± 0.1 ppm, or air, in inhalation chambers for 6 hours per day, 5 days per week for 12 weeks.[32, 35] Body weight and food consumption were monitored throughout the exposure regimen. Animals were sacrificed either immediately following 12 weeks of exposure or following an additional 9 months of housing (±WD feeding). At sacrifice, mice were fasted for 4 hours and anesthetized with ketamine/xylazine (100/15 mg/kg, i.p.). Blood was collected from the vena cava, and citrated plasma was stored at −80°C. Liver tissue was snap-frozen in liquid nitrogen, embedded in frozen specimen medium (Sakura Finetek, Torrance, CA), or was fixed in 10% neutral buffered formalin.

Additional methods are presented in the Supplemental information.

## Results

### VC increases WD-induced tumor formation

Recent studies by this group determined the effects of VC inhalation at levels below the current safety regulations (<1 ppm) in the context of NAFLD to mimic human exposure and identify potential mechanisms of VC induced liver injury.[32, 35] In that model, VC exposure caused no overt liver injury in mice fed CD. In mice fed WD, VC significantly increased liver damage as shown by elevated serum transaminases and steatosis (Figures 1B-D). VC caused metabolic dyshomeostasis and enhanced WD-induced oxidative and endoplasmic reticulum stress.[36] Here, following the 12-week VC exposure, mice were continued to be fed WD (or CD) for 9 months (timeline, Figure 1A). While CD livers were normal, pericentral and midzonal steatosis which was predominantly macro-vesicular (Figure 1D). WD feeding for 1 year caused almost panzonal steatosis (macro- and micro-vesicular), while VC+WD caused panzonal macro- and micro-vesicular steatosis and well circumscribed neoplastic foci with no or modest steatosis (Figure 1D). In the WD group, 20% of the livers showed the presence of neoplastic foci. In contrast, in the VC+WD group, 100% of the livers showed tumors, which were validated to be HCC (tumor outlined, H&E stain and glutamine synthetase IHC, Figures 1B, 1D and 2A).

**Fig. 1:**
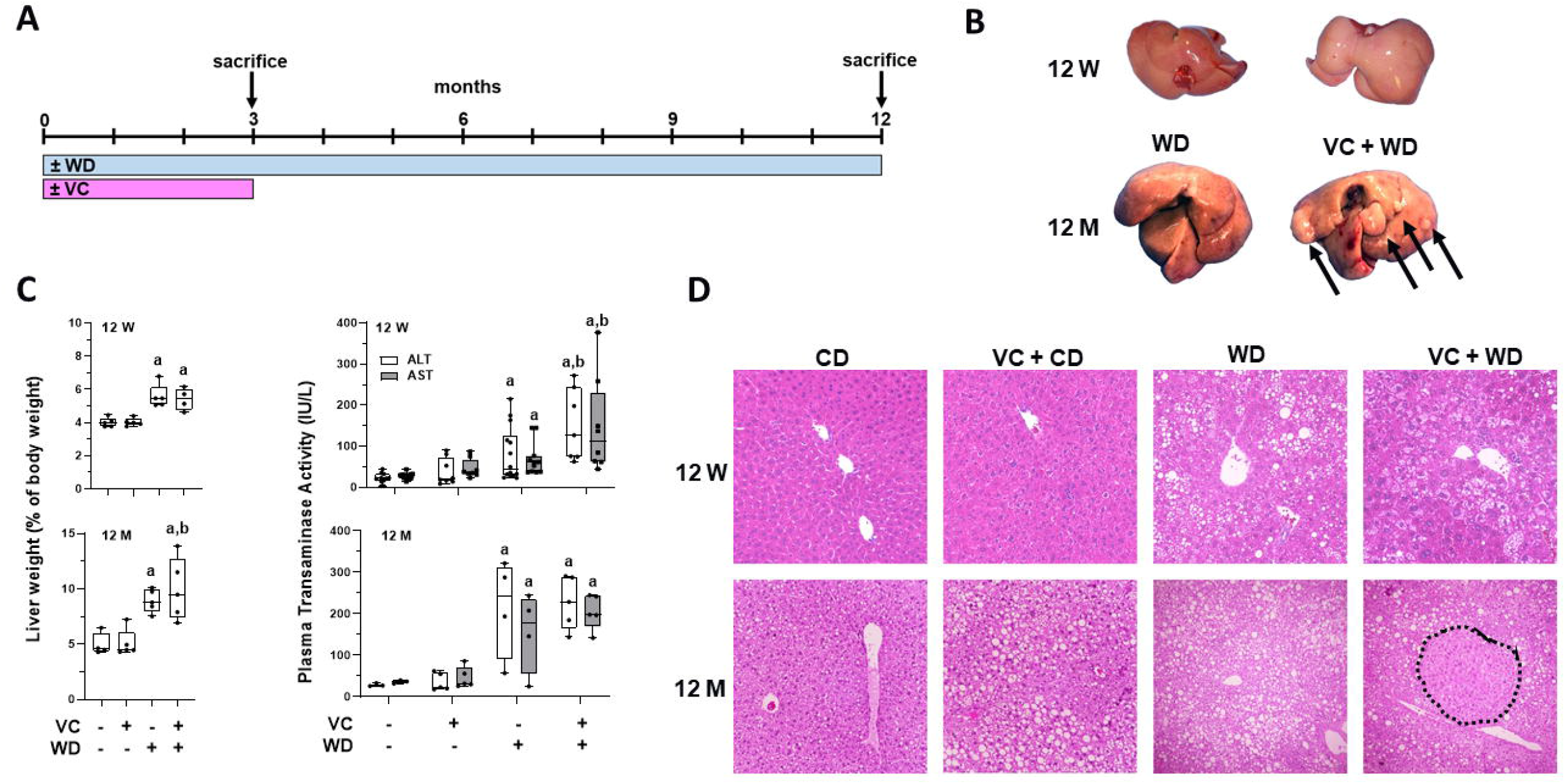
VC enhances WD-induced tumorigenesis. **A:** Timeline of exposure, feeding and sacrifice regimen. **B:** Gross liver comparison at 12 weeks (12 W) and 12 months (12 M). Visible tumors at 12 M in the VC + WD group only (denoted via arrows). **C:** VC significantly increased LW:BW ratio, tumor number/size in the WD group only at the 12-month timepoint. **D:** VC enhances WD-induced elevation in transaminases (ALT and AST) after 12 W. There was no difference in transaminase levels 9 months after cessation of VC exposure in the CD or WD group. **E:** H&E stain at 12 W and 12 M; HCC outlined. a, p<0.05 compared to control; b, p<0.05 compared to no VC. n=5 per group.

**Fig. 2:**
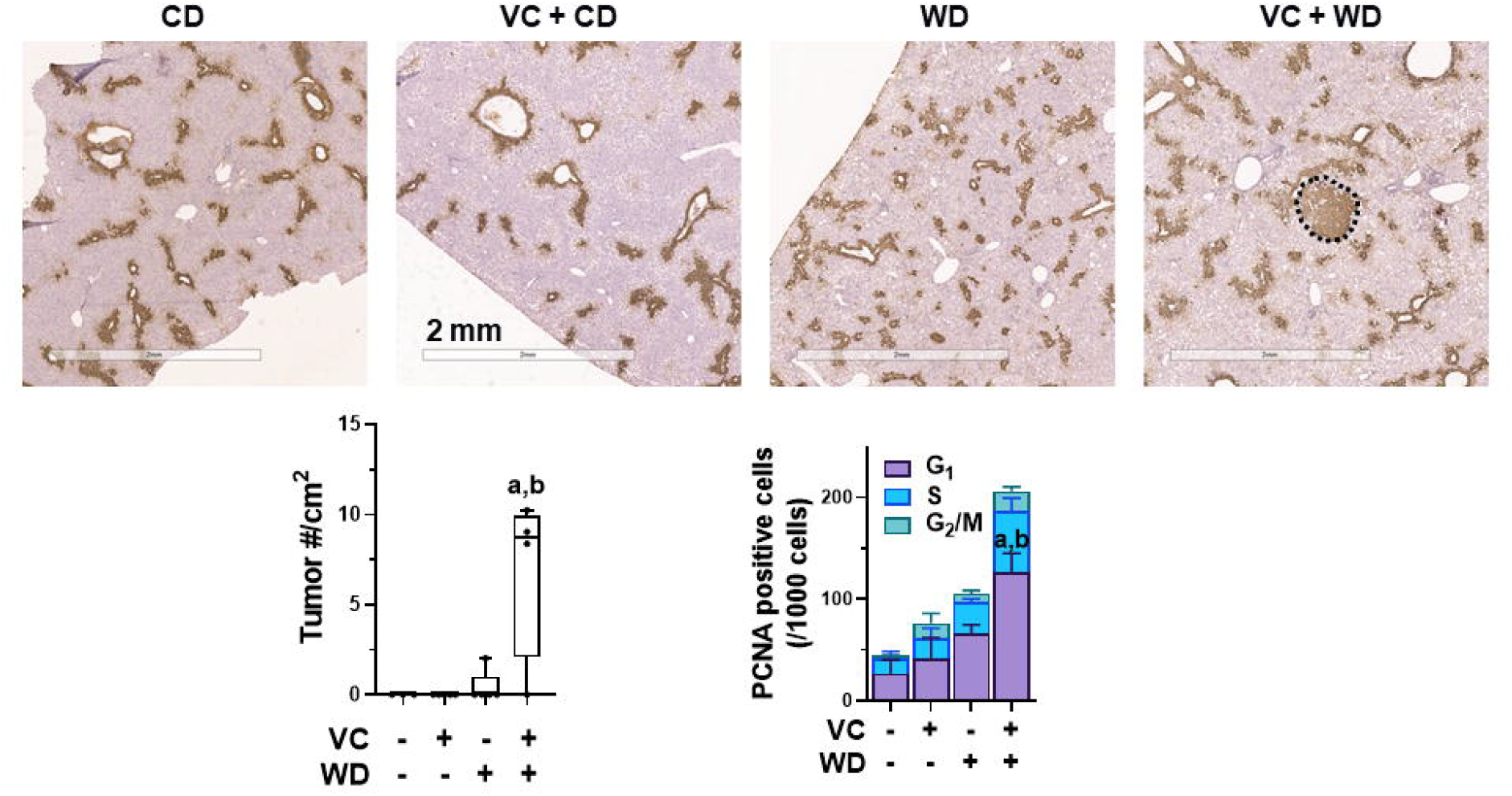
VC enhances WD-induced tumor number and proliferation. **A:** Glutamine synthetase IHC; HCC outlined. **B:** VC significantly increased tumor number/cm^2^ and number of PCNA+ cells in the WD group. a, p<0.05 compared to control; b, p<0.05 compared to no VC. n=5 per group.

Tumor burden is depicted in liver weight (LW):body weight (BW) ratios and tumor number/cm^2^ (Figures 1C and 2). The tumors were visible as well-circumscribed lesions with Ki67^+^ (Supplemental Figure 1) and PCNA^+^ cells (Figure 2). Pathologic assessment demonstrated tumors ranging from moderately to poorly differentiated HCC with clear basophilic cytoplasm, moderate to high nuclear atypia. While there was no fibrosis in the absence of WD, there was evidence of fibrosis in WD, which was more extensive in VC+WD (Sirius red staining; Supplemental Figure 2). Thus, 12 weeks transient exposure to VC at the beginning of WD feeding clearly exacerbated the normal tissue damage and led to development of HCC.

### The enhancement of HCC by VC exposure leads to a unique transcriptomic phenotype

To identify potential mechanisms by which VC exposure enhanced HCC development, transcriptomic analysis (RNA-Seq) was performed. VC exposure (in contrast to air controls) significantly changed the expression of several genes at 12 months (up: 201, down: 168; Figure 3), a robust increase in changes compared to the 12-week time point (up: 70, down: 60; Supplemental Figure 3). Based on these differentially expressed genes (DEGs), GO, KEGG and JASPAR pathway analyses were performed, demonstrating 62 significantly changed pathways, including several involved in metabolic processes such as lipid metabolism that were robustly induced at 12 months (Figure 3). More specifically, these included GO terms for fatty acid metabolic process, cholesterol metabolic process, and sterol metabolic process. In addition, Ingenuity Pathway Analysis (IPA) was performed on the DEGs. Employing the IPA Upstream Regulator analytic, PPARα signaling (Z-score=-2.0; p=8.5×10^−13^) was predicted to be downregulated in WD-fed mice exposed to VC vs. air. Despite the robust increase in cholesterol metabolic processes, there was no difference in oil red-o (ORO) positive staining and cholesterol or cholesterol esters between WD and WD with transient VC exposure (Supplemental Figure 2).

**Fig. 3:**
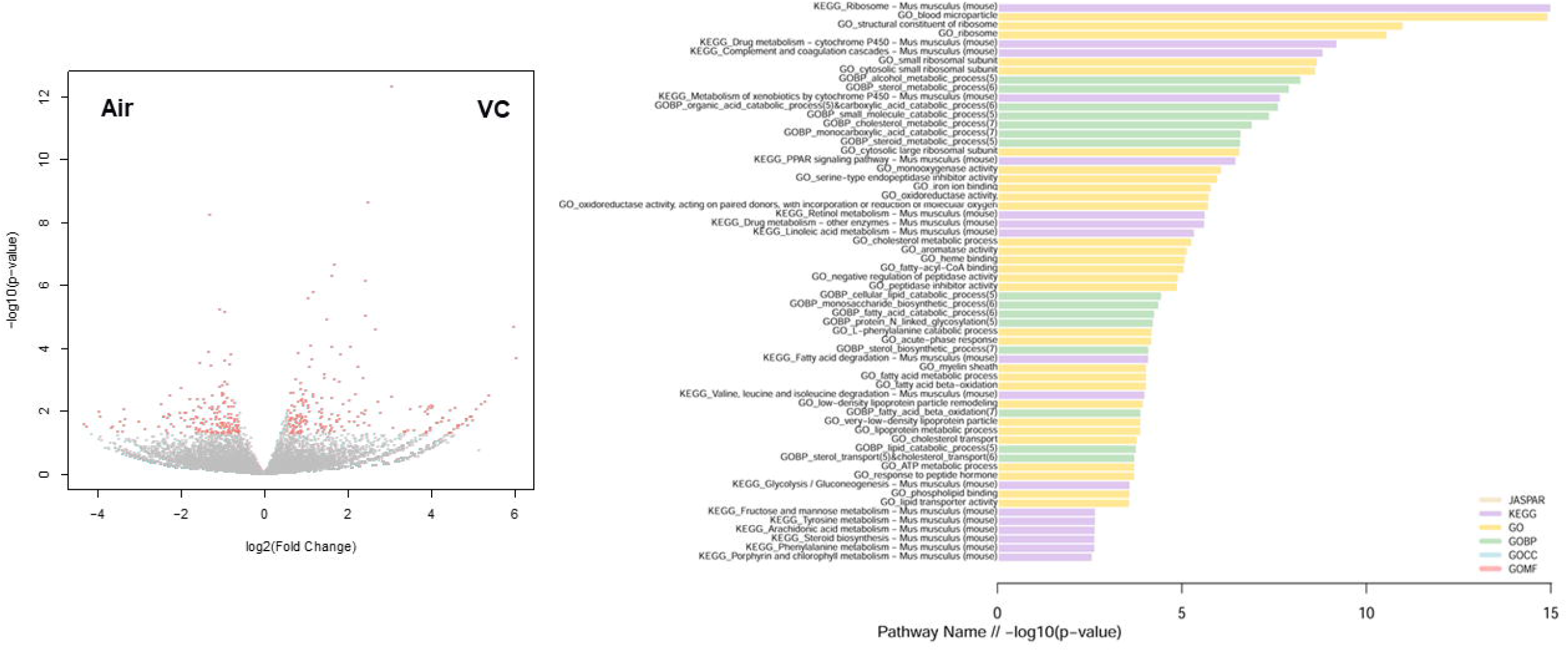
RNA-Seq analysis. n=4 per group. Left panel: volcano plot comparing the expression pattern in mice exposed to VC (versus air) at the 12-month timepoint. Each dot represents a gene, with red dots highlighting significantly up- or down-regulated genes. Right panel: top KEGG, GO, and GO BP terms identified changed by pathway enrichment analysis.

In addition to the changes in metabolic processes, VC also caused major changes in ribosomal processes and RNA processing at the 12-month time point (Figure 3). These changes were very similar to changes over time (WD+VC 12 months vs. 12 weeks; Figure 4). When the expression pattern caused by VC exposure was analyzed (up: 213, down: 267, 26 significant pathways, Figure 4B), employing the IPA Upstream Regulator analytic, dramatic changes in key regulators of metabolic pathways and ribosomal processes, including MLXIPL (up; carbohydrate-responsive element binding protein, ChREBP), MYCN (up; N-Myc: regulates ribosome biogenesis and mitochondrial gene expression programs), ACOX (down; involved in β-oxidation) and LARP1 (down; involved in ribosomal biogenesis) (Figure 4C). This was also reflected in the ‘top significant pathways’ (Figure 4D, Supplemental Table 2), highlighting the EIF2 signaling pathway, which is involved in translation, initiation, and assembly of the ribosomal subunits.

**Fig. 4:**
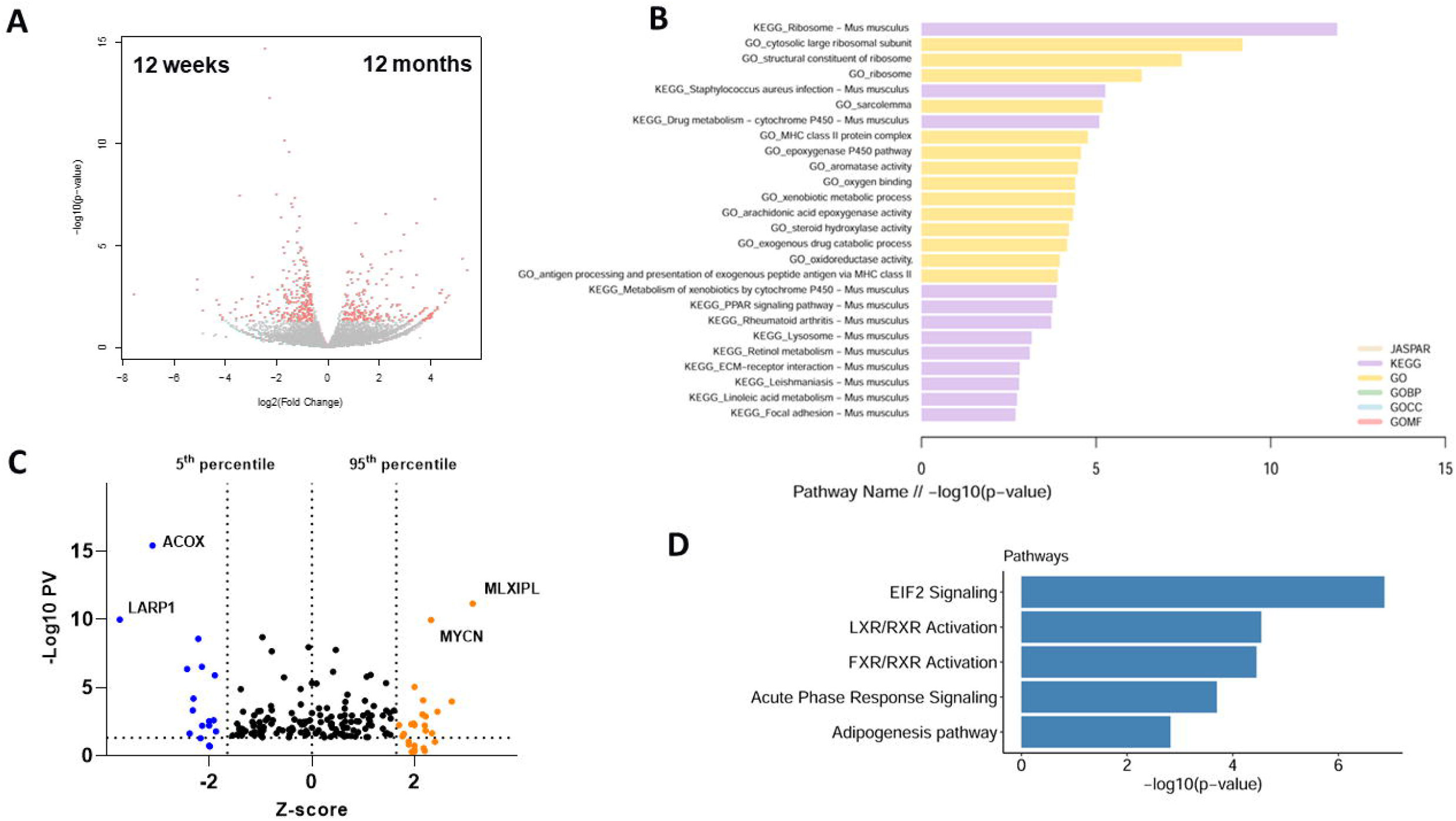
RNA-Seq pathway analysis. n=4 per group. **A:** volcano plot comparing the expression patterns at 12-months vs. 12-weeks. Each dot represents a gene, with red dots highlighting significantly up- or down-regulated genes. **B:** top pathways identified changed by pathway enrichment analysis. **C:** volcano plot based on IPA upstream regulator analysis, with each dot representing the transcription factors on livers of mice exposed to VC+WD at 12 weeks vs. 12 months. The p-value for significance is plotted as a function of the Z-score. **D:** Significant pathways detected by IPA. **ACOX:** first enzyme in β-oxidation pathway; decrease associated with poor outcome in HCC. **LARP1:** RNA-binding protein; ribosome biogenesis. **MLXIPL:** (ChREBP) promotes HCC proliferation. **MYCN:** (N-Myc) proto-oncogene; associated with HCC. **EIF2:** stress-related master controller of RNA processing.

### Biomarkers in mouse model provide insight in human HCC cohort

The RNA-Seq results provide weight-of-evidence that VC can enhance the development of HCC, as demonstrated by ‘top significant diseases and disorders’ (Supplemental Figure 4), via mechanisms likely involving altered ribosomal processes (Figure 4 and Supplemental Figure 4). However, whether these results have relevance to human HCC remained unclear. To address this, all changed genes that are involved in ribosomal processes (KEGG: 03010) were applied to human ortholog[37] expression in the Liver Hepatocellular Carcinoma (LIHC) sequencing results in the Cancer Genome Atlas (TCGA). These signature genes clustered all the TCGA tumor samples into 5 hierarchical clusters (Figure 5A). Interestingly, a sub-cluster (green cluster, Figure 5A) highly correlated with the murine expression pattern. To determine if the tumors occurring in WD-fed mice exposed to transient VC share molecular signatures with HCC patients, the degree of similarity between the human differential expression analysis of HCC cases and the VC mouse model were assessed. First, differentially expressed genes (DEGs) from the TCGA cohort comparing normal vs tumor samples and from the mouse model were selected. Next, mouse DEGs were mapped to their human orthologs according to the MGI mouse-human ortholog database (http://www.informatics.jax.org/). Common up- and down-regulated genes in the two cohorts were shown in Figure 5B. This analysis revealed a significantly high correlation between the sets of changes in the two cohorts, with a Spearman’s correlation of 0.71 with a p-value of 2.2×10^−16^ and with a Pearson correlation of 0.681 with a p-value of 9.741×10^−13^. These data indicate that the key transcription changes observed in the mouse model of VC exposure have strong expression concordance with a subset of human HCC.

**Fig. 5:**
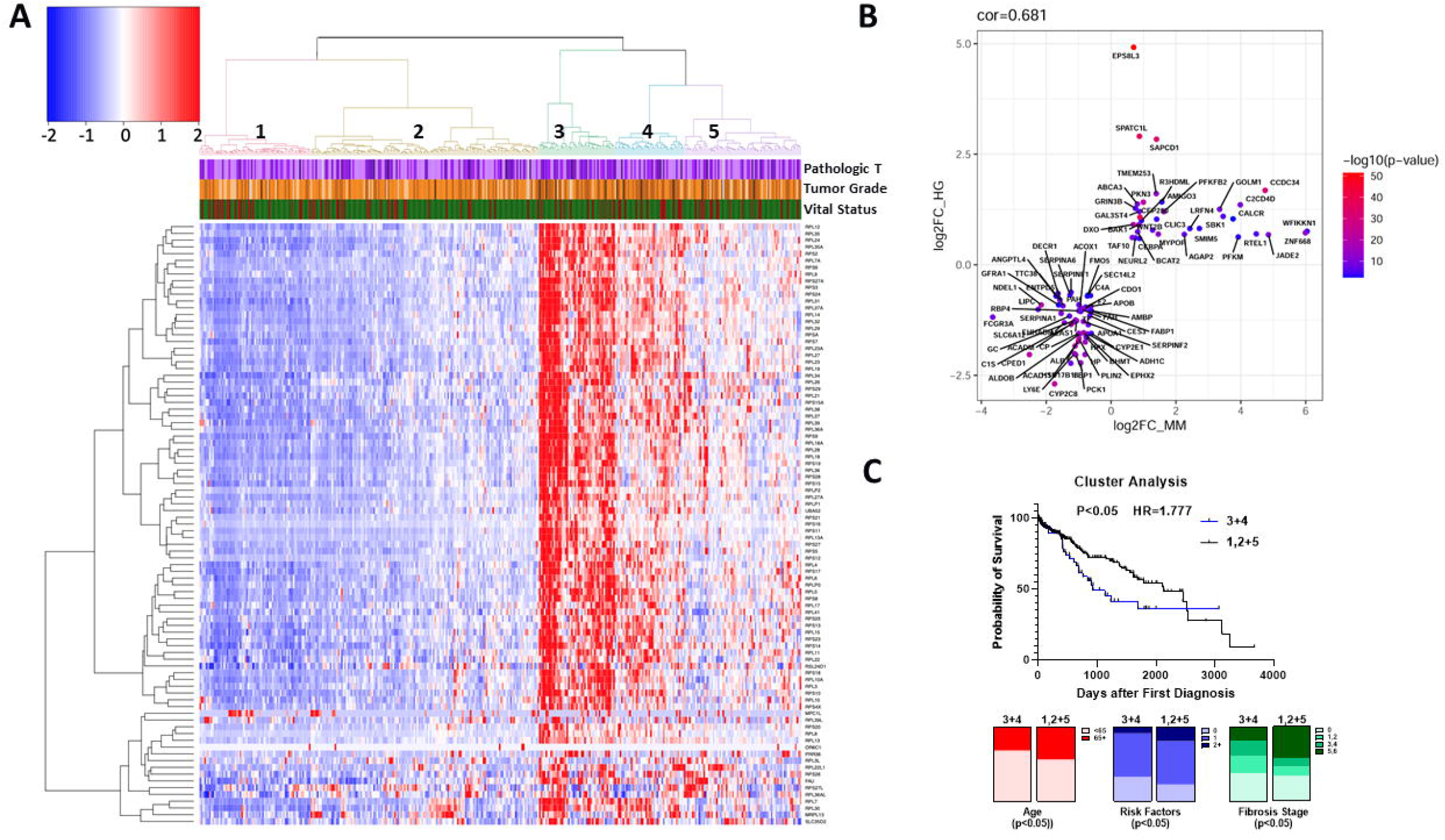
TCGA data analysis. A unique gene signature of VC-induced tumors correlates with a subset of human HCC. **A:** Heatmap of the human TCGA samples divided based on the 89 genes involved in ribosomal pathways and changed by VC exposure after 12 months (see Figures 3 and 4). Cancer grade (Pathologic T and Tumor Grade) and mortality (vitality; green=alive) are also shown for individual HCC cases. **B:** A differential gene expression analysis of the human HCC subset comparing normal liver vs. tumor was performed and the extent to which these gene changes were recapitulated in the mouse model was assessed. Common up- and down-regulated in both human and mouse models are highly correlated with a Spearman correlation of 0.71 with a p-value of 2.2×10^−16^ and with a Pearson correlation of 0.681 with a p-value of 9.741×10^−13^.

The clinical variables from the TCGA database associated with the 5 hierarchical clusters were then analyzed. The 2 clusters associated with an elevated expression pattern of ribosomal proteins cluster 3 (green) and cluster 4 (blue) were compared with the other clusters. Univariate Chi-square analysis indicated that patients in Clusters 3 and 4 were more evenly distributed by sex, younger, with fewer primary risk factors for HCC (e.g., alcohol, NAFLD, HCV, etc.), and had less severe CLD (Ishak scale; Figure 5C and Table 1). In fact, the number of patients with cirrhosis (Ishak 5 and 6) was ~50% lower in clusters 3 and 4 compared with the other clusters. Although these factors would be expected to correlate with better survival, total mortality tended to be ~40% higher (p=0.0767), and Kaplan-Meier assessment of the predicted survival indicated that these groups actually had a steeper rate of mortality than the other 3 clusters by unadjusted Cox Proportional Hazard analysis (HR=1.65, 95% CI 1.04-2.63, p=0.035). This Hazard Ratio was not increased by multivariate analysis of known risk factors for HCC mortality (e.g., age, pathologic T, tumor grade, fibrosis score and number of HCC risk factors; not shown).

### The enhancement of HCC by VC exposure leads to a unique epitranscriptomic phenotype

Emerging evidence indicates that mRNA sequence is by no means the sole regulator of post-transcriptional gene regulation. This process is also regulated by secondary mRNA structure (i.e., folding) and chemical modifications of RNA bases, which is similar in concept to DNA (i.e., methylation, acetylation, etc.). These interdependent non-sequence features of mRNA (i.e., epitranscriptomics) exert direct control over the transcriptome and thereby influence many aspects of cell function independent of mRNA abundance. Importantly, RNA modifications may also alter RNA-protein interactions during gene expression. Therefore, an LC-MS/MS analysis of total hepatic nucleoside and nucleotides was performed on samples from the 12-week (Supplemental Figure 5 and Supplemental Table 3) and the 12-month time points (Figure 6). An analysis of changes over time (WD+VC, 12 months vs. 12 weeks, Figure 6 and Supplemental Table 3) was also performed. VC induced significant changes to the modification pattern (Figure 6) that have been associated with cancer (e.g., m3C, m6A, m1A, and Gm), but also with mitochondrial dysfunction and metabolic disease (e.g., m4C, m6A, and m1A).

**Fig. 6:**
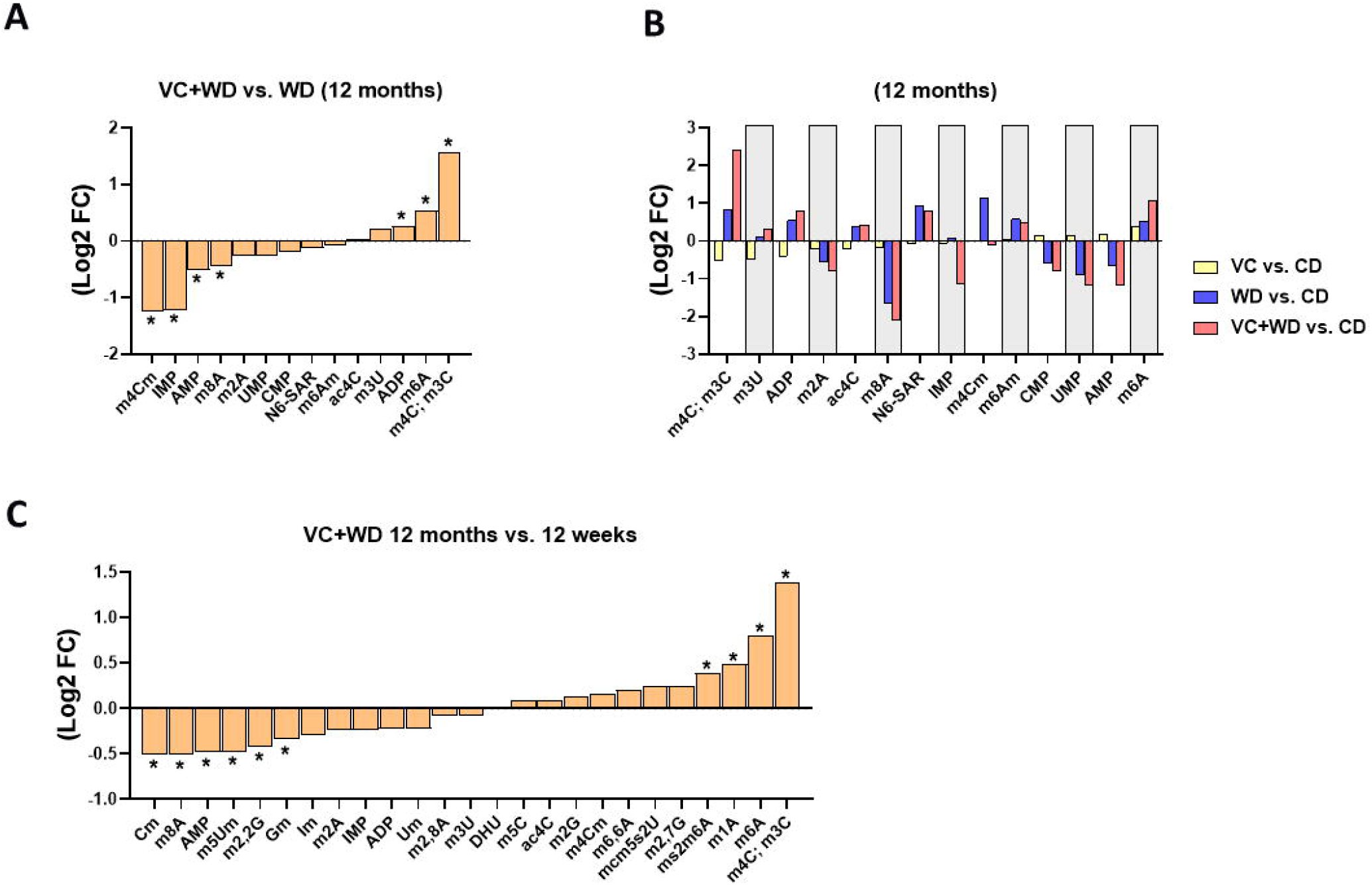
VC changes the pattern of total hepatic nucleosides/nucleotides. LC-MS/MS epitranscriptomic analysis of total hepatic nucleosides and nucleotides at 12 months and 12 weeks vs. at 12 months. *, p<0.05 (FDR) log2 fold change (FC). n=5 per group.

## Discussion

HCC occurs predominantly in the context of CLD where cycles of hepatocyte injury, inflammation and regeneration trigger survival and proliferation of hepatocytes harboring oncogenic mutations. Therefore, the incidence of HCC has not surprisingly paralleled the increase in CLD. Although advanced therapeutic options have recently improved survival for most cancers, the low survival rate for HCC has not changed for decades, although the advent of immunotherapy is showing promise in a subset of HCC cases. These factors taken together explain why HCC-related deaths continue to rise.[38] It has been demonstrated that multiple parallel hits drive not only CLD pathogenesis, but also the development of HCC on the background of CLD.[39] Although environmental exposure is increasingly identified as a potential risk factor for CLD,[40] few studies have investigated the impact of environmental exposure on the risk of developing HCC. The purpose of this study is to begin to address this gap in our knowledge.

High occupational VC exposure is well-known to be associated with the development of hemangiosarcoma and HCC,[41–43] which were largely eradicated with a lowering of exposure limits in the 1970’s. However, these safety guidelines are potentially outdated, considering the significant increase in average BMI and NAFLD/NASH across the global population in the subsequent years.[44, 45] Therefore, it has been unclear if the interaction between VC exposure and underlying liver disease enhances the risk for the development of liver cancer at exposure levels lower than the current safety limits. The primary goal of the current study was to study the process underlying the development of HCC with a newly developed animal model of exposure to low levels of VC. The present study’s findings demonstrated that transient (12 weeks) VC exposure accelerates and enhances tumorigenesis caused by experimental NAFLD in mice. This enhancement of HCC appears not to be simply via a worsening of the underlying CLD, but rather a direct effect on HCC pathogenesis, as most indices of liver damage were not significantly altered between the air- and VC-exposed cohorts (Figures 1–2).

Pathway Analysis showed that VC exposure altered several metabolic processes involved in metabolism (Figure 3), including GO terms for fatty acid metabolic process (GO:0006631; p=9.3×10^−5^), cholesterol metabolic process (GO:0008203; p=1.3×10^−7^), and sterol metabolic process (GO:0016126; p=1.3×10^−8^). When the expression pattern caused by VC exposure was analyzed, employing the IPA Upstream Regulator analytic, dramatic changes in key regulators of lipid synthesis, including ChREBP (MLXIPL; up), ACOX1, and PPARα (down) were observed (Figure 4). This finding is in-line with previous work showing that VC exposure alters metabolism in precancerous models.[32, 36, 46] Although altered metabolism is an established key factor in the development of CLD, more recent work has identified that it also may contribute to cancer biology and the development of HCC.[47–51] Moreover, ChREBP activation has been previously shown to promote HCC proliferation and has been associated with poor outcome in patients with HCC.[52]

The key pathways identified by pathway enrichment analyses were associated with increases in ribosomal processes, including ribosomal biogenesis, structural constitution, and RNA processing. Moreover, IPA Upstream Regulator analytic identified strong induction of the EIF2 signaling pathway, a major mediator of translation, initiation, and assembly of ribosomal subunits. This finding is in-line with previous work in humans, which has identified t-RNA aminoacylation, the initial step in translation initiation, as the top canonical pathway affected by occupational VC exposure.[53] It is well established that altered ribosomal processes and riboslome biogenesis play an important role in cancer, as any quantitative change in ribosome concentration may impact translation patterns and favor expression of specific mRNAs to the detriment of others.[54] In cancer cells, disruption of ribosome biogenesis and protein synthesis is associated with altered expression of key genes encoding translation initiation factors and proto-oncogenes such as MYC.[55] Indeed, here IPA Upstream Regulator analytic revealed that transient VC not only induced major changes in expression of, but also strongly induced, n-MYC (MYCN). Moreover, MYC and ChREBP transcription factors have been shown to cooperatively regulate normal and neoplastic hepatocyte proliferation in mice.[56] These novel findings were not observed in previous studies of VC exposure in precancerous models and were not associated at earlier (12 weeks) sacrifices in this study (Supplemental Figure 3). These results suggest that the alteration of ribosomal processes is not directly related to the liver injury caused by VC exposure, per se.

The mechanisms by which ribosomal processes were induced under these conditions remain unclear. This may be a direct result of the metabolic changes caused by VC exposure. For example, previous studies have shown that activated ChREBP pathways can lead to upregulation of ribosomal processes.[56] However, recent work has indicated that direct modifications of ribosomes and/or RNAs may influence these processes. This is especially important as about 2/3 of the ribosome is comprised of RNA. Cellular RNAs are subject to multiple chemical modifications, and the study of these chemical modifications (i.e. the epitranscriptomics) is a growing research interest in disease and dysfunction.[57–59] The work to date indicates that epitranscriptomic alterations play important roles in regulating RNA metabolism, translation, localization, stability, turnover, as well as in binding to proteins or other RNAs, which ultimately affects metabolism, disease progression, and carcinogenesis. While qualitative modifications can dictate oncogenic potential, the knowledge about specific modifications and their role in carcinogenesis is still limited. However, N6-methyladenosine (m^6^A), regarded as the most abundant internal modification, has been associated with poor prognosis in HCC.[60, 61] Moreover, m^6^A methylation sites have been shown to be enriched in processes associated with lipid metabolism.[61] Importantly, VC strongly increased m^6^A abundance at the 12-month time point, suggesting a potential role for m^6^A in the metabolic and tumorigenic changes caused by VC. Moreover, it has been shown that together with the EIF2 signaling pathway, which was strongly induced here, mRNA methylation in the form of m^6^A reconfigures the cellular adaptation to stress conditions.[62]

VC also increased total m^3^C and m^4^C. The 3-methylcytosine (m^3^C) modification is present in both tRNA and mRNA and displays diverse roles in developing diseases through the regulation of tRNA fate. It has been shown recently that methylation in tRNA enhanced the proliferative activity of hepatocellular carcinoma (HCC).[63, 64] Currently not much information is known about the 4-methylcytosine (m^4^C) modification and its impact on human disease; however m^4^C on mitochondrial rRNA has been associated with mitochondrial function. It should be noted that one limitation of this study is that while LC-MS/MS analysis is highly sensitive and quantitative, it cannot distinguish between m^3^C and m^4^C nor determine which RNA sites are modified. Furthermore, here changes to whole liver nucleoside/nucleotide modifications were measured. Therefore, future studies will need to be performed to determine the location of the modification and RNA species.

The increase in ribosomal processes may also have an impact on HCC development and outcome. Two clusters of human HCC were identified from the TCGA database that were enriched for increases in expression of genes associated with ribosomal processes. Interestingly, HCC in these groups more frequently occurred on the background of less severe CLD and with fewer known risk factors. This group also had a higher Hazard Ratio for mortality. This administrative and retrospective analysis does not allow identification of other causes that may drive HCC. However, there was strong concordance in the overall expression patterns of this human HCC and the murine orthologs from WD-fed mice exposed to VC. It is therefore distinctly possible that unidentified environmental exposure may be a hidden factor that contributed to the development of HCC in this cohort. Since several environmental chemicals like VC may impart similar effects on the target organ,[29, 40, 65] this potential environmental exposure may be representative of more than VC per se, and by extension, may represent exposures that also cause epitranscriptomic changes. These results bolster the need for more comprehensive documentation of exposure (i.e., the “exposome”).[66]

This is the first study to demonstrate VC-induced tumorigenesis at concentrations that are currently considered safe. In summary, the data indicate that VC sensitizes the liver to other stressors (e.g., WD), involving dysregulation of metabolic pathways, an enrichment of genes for ribosomal processes, and changes to the epitranscriptomic pattern. These changes result in enhanced tumorigenesis and correlate with a subset of human HCC that tended to be younger with fewer primary risk factors for HCC. These data raise concerns about potential for overlap between hypercaloric diets and exposure to lower concentrations of VC, as well as the health implications of this co-exposure for humans that may be underappreciated with our current knowledgebase. It also emphasizes that current safety restrictions may be insufficient to account for other factors that can influence hepatotoxicity.

---

| **Measure** | **Category** | **Clusters**<br>**3+4**<br>**(n=90)** | **Clusters**<br>**1,2+5**<br>**(n=284)** | **p-**<br>**value** |
| --- | --- | --- | --- | --- |
| **Age at first**<br>**Diagnosis** | **Years (±SD)**<br>**<65 (%)**<br>**65+ (%)** | 57.6±13.9<br>68.9<br>31.1 | 60.0±13.3<br>56.5<br>43.5 | 0.1237<br>0.0376 |
| **Sex** | **Male (%)**<br>**Female (%)** | 60.0<br>40.0 | 70.0<br>30.0 | 0.0376 |
| **Race** | **White (%)**<br>**Black (%)**<br>**Asian (%)**<br>**Am. Indian (%)** | 41.1<br>3.3<br>52.2<br>1.1 | 52.3<br>4.9<br>39.9<br>0.4 | 0.1549 |
| **Risk Factors** | **None (%)**<br>**1 (%)**<br>**2+ (%)** | 33.3<br>58.3<br>8.3 | 23.2<br>57.9<br>18.9 | 0.0324 |
| **Fibrosis Score**<br>**Ishak Scale** | **0 (%)**<br>**1,2 (%)**<br>**3,4 (%)**<br>**5,6 (%)** | 38.5<br>23.1<br>20.5<br>17.5 | 34.1<br>12.5<br>11.4<br>42.0 | 0.0220 |
| **Tumor Stage** | **I (%)**<br>**II (%)**<br>**III (%)**<br>**IV (%)** | 40.7<br>26.7<br>30.2<br>2.3 | 52.3<br>24.4<br>22.3<br>1.1 | 0.2383 |
| **Tumor Grade**<br>**pT** | **T1 (%)**<br>**T2 (%)**<br>**T3 (%)**<br>**T4 (%)** | 40.0<br>27.8<br>27.8<br>4.4 | 52.3<br>24.9<br>19.6<br>3.2 | 0.1917 |
| **Deaths** | **(%)** | 34.25 | 23.97 | 0.0767 |
| **Unadjusted HR** | **HR (95% CI)** | 1.65 (1.04-2.63) |  | 0.035 |

