## supplemental material for "Western diet unmasks transient low-level vinyl chloride exposure-induced tumorigenesis; potential role of the (epi-)transcriptome"

**SUPPLEMENT**

A. **Supplemental Methods:**

**Biochemical Analyses.** Plasma levels of ALT & AST were determined using standard kits (Thermo Fisher Scientific, Middletown, VA).

**Histology.** Paraffin-embedded sections of the liver were stained with hematoxylin & eosin (H&E) to assess the overall hepatic structure. Cellular proliferation was visualized via Ki-67 immunohistochemistry using the rabbit anti-mouse primary antibody (1:100). CD31 immunohistochemistry was done for angiogenesis using the rabbit anti-mouse primary antibody (1:100). Oxidative stress was determined by visualizing the 4-hydroxynonenal (4-HNE) adducts via immunohistochemistry using the rabbit anti-mouse primary antibody (1:500, Alpha Diagnostics, San Antonio, TX). Frozen samples kept in OCT compound were cut and used for Oil Red O staining (for neutral lipids) and fibrin immunostaining. Sections were stained with Oil Red O (Sigma O-9755) solution for 10-12 minutes and then rinsed in distilled water before staining with hematoxylin and mounting with aqueous media. Fibrin was visualized using polyclonal rabbit anti-human fibrinogen antibody (1:1000 Dako, Carpinteria, CA). Glutamine synthetase was visualized by indirect immunohistochemistry (1:50, sc-74430, Santa Cruz). Collagen formation was assessed by visualization of Sirius Red (SR) stained liver sections. Image analysis was performed using Metamorph Image Analysis Software (Molecular devices, Sunnyvale, CA) and expressed as positive staining in % of microscope field.[1] The sample size for all the immunohistochemistry and staining was between 4-5. For analysis, 10 randomly selected images were taken from each sample and analyzed using the Metamorph Image Analysis Software.

**RNA-Seq and data analysis.** Total RNA was extracted using Trizol and treated with DNAse1. Ribosomal RNA was removed from the samples using RIBO-ZeroTM Magnetic kit (Epicentre, Madison, WI).[2, 3] RNA was reverse-transcribed to cDNA and amplified using TruSeq™ RNA Sample Prep Kit v2 from Illumina, Inc (San Diego, CA). The cDNA library was similarly sequenced in Illumina HiSeq2500 sequencer.[4] Quality control on raw RNA-Seq data was performed using the tool FastQC[5] and low-quality reads and adapter sequences were filtered out using the tool Trimmomatic.[6] The surviving reads were then aligned to the mouse reference genome mm10 by STAR aligner, and the genes were quantified using the tool HTSeq.[7] Then differential expression analyses, using R package DESeq2,[8] were performed to compare the pairwise experimental groups. Differentially expressed genes (DEGs) were defined by FDR=5% and fold-change equal to or greater than 1.5. These DEGs were further used as input for ingenuity pathway analysis (IPA)[9, 10] to detect significantly-enriched pathways and common upstream regulators. Pathway analyses on Gene Ontology (GO),[11] Kyoto Encyclopedia of Genes and Genomes (KEGG)[12-14] and JASPAR[15] databases were also performed using the DEGs. Significant pathways were defined by FDR=5%.

**The Cancer Genome Atlas (TCGA) data mining.**

To explore the TCGA database, gene expression profiles for 374 tumors and 50 normal (or adjacent tumor) tissues with HCC and RNA sequencing data were collected from Genomic Data Commons (GDC) Data Portal (https://portal.gdc.cancer.gov/). Differential expression analysis comparing normal and tumor samples were performed by R package DESeq2.[8] The potential association between RNA expression patterns and TCGA clinical features of HCC patients was compared, which included age of first diagnosis, gender, race, risk factors, fibrosis score (Ishak scale), tumor stage, tumor grade and vital status. The frequency of missing data at baseline was 0% for sex, 0.2% for Age, 2.4% for race, 5.1% for Risk Factors, 42.5% for fibrosis score, 6.1% for tumor stage, 0.8%. Vital status for 8.8% patients was lost to follow-up.

In order to compare the mouse study with human TCGA cohort, DEGs comparing the experimental groups in the mouse model were recruited as biomarkers. The mouse genes were first mapped to human homology genes via the MGI database.[16] Then these markers were applied to the TCGA[17] liver hepatocellular carcinoma (LIHC) cohort. Hierarchical clustering was then performed on the tumor cases to group all the HCC patients into subtypes based to the mouse signatures selected in our study. The resulting marker genes were grouped into clusters as well. Kaplan-Meier survival curves were plotted on the two groups clustered by these gene markers and p-value was calculated by the log rank test. To further explore the consistency between mouse and human studies, common differentially expressed genes in both our mouse model and the TCGA cohort were detected. Pearson correlation and Spearman’s correlation were calculated on these common DEGs.

**Epitranscriptomic analysis.** Total RNA was extracted from whole liver. RNA extracts were denatured at 90 °C for 3 min and then chilled in an ice-water for 3 min. To detect modifications of RNA on nucleosides level, 15 µg RNAs were first mixed with 240 units nuclease S1 and 2 μL of reaction buffer with pH= 4.5 (500 mM sodium acetate, 2.8 M NaCl, and 45 mM ZnSO_4_). The mixture was incubated at 37^o^C for 4 h. After removing nuclease S1 using a microcon centrifugal filter (Microcon YM-10), the pH of the solution was adjusted to basic using 2.5 µL of a mixture with pH = 9.3 (1 mM MgCl_2_, 0.1 mM ZnCl_2_, 1 mM spermidine, 50 mM Tris-HCl) and 0.5 units of phosphodiesterase I was added. The solution was incubated at 37 ^o^C overnight, and phosphodiesterase I was then removed using Microcon YM-10. Finally, 3 units of antarctic phosphatase were added. The solution was incubated at 37 ^o^C for 4 h and antarctic phosphatase was removed using Microcon YM-3.

All samples were analyzed randomly on a Thermo Q Exactive HF Hybrid Quadrupole-Orbitrap Mass Spectrometer coupled with a Thermo DIONEX UltiMate 3000 HPLC system (Thermo Fisher Scientific, Waltham, MA, USA) using the method developed previously.[18] To obtain full MS data, every sample was analyzed by LC-MS in positive mode (+). For metabolite identification, one pooled sample for each group was analyzed by LC-MS/MS in positive mode to acquire MS/MS spectra at three collision energies (20, 40, and 60 eV).

For LC-MS data analysis, XCMS software was used for spectrum deconvolution,[19] and MetSign software was used for cross-sample peak list alignment, normalization, and statistical analysis[20-22]. To identify nucleosides, the LC-MS/MS data of the pooled samples were matched to the MS/MS spectra of 94 nucleoside standards recorded in Compound Discoverer (v3.2, Thermo Scientific) that contains parent ion m/z value, retention time, and MS/MS spectra. Chemspider and mzCloud were used for the identification of unknown compounds.[23] The threshold for the spectral similarity of the MS/MS spectra of a nucleoside standard and a spectrum of the pooled sample was set as ≥ 40 with a maximum score of 100, while the thresholds of the retention time difference and m/z variation window were set as ≤ 0.15 min and ≤ 5 ppm, respectively.

**Statistics.** Summary data represent means ± SEM (n = 4-5). ANOVA with Bonferroni's post-hoc test or the Mann-Whitney rank sum test was used for the determination of statistical significance among treatment groups, as appropriate. A p-value < 0.05 was selected before the study as the level of significance.

**Data Availability:** Transcriptome sequencing data generated in this study were deposited into NCBI Gene Expression Omnibus (GEO), with accession ID GSE197038. Raw sequencing reads and gene quantification profile can be downloaded at <https://www.ncbi.nlm.nih.gov/geo/query/acc.cgi?acc=GSE197038>.

B. **Tables:**

**Supplemental Table 1-** **Chemical assays used in this study**

| **Category** | **Name** | **Supplier** | **Cat No.** |
| --- | --- | --- | --- |
| *Chemical assays* | AST assay kit | Thermofisher | TR70121 |
|  | ALT assay kit | Thermofisher | TR71121 |

**Supplemental Table 2: Transcriptomic analysis**

| **Top significant pathways** | | | | | | | | | | | | |
| --- | --- | --- | --- | --- | --- | --- | --- | --- | --- | --- | --- | --- |
| **Ingenuity Canonical Pathways** | | **-log(p-value)** | | **Molecules** | | | | | | | | |
| EIF2 Signaling | | 6.87E+00 | | DDIT3,FAU,PPP1CA,RPL13,RPL13A,RPL14,RPL19,RPL21,RPL37A,RPL4,RPL6,RPL8,RPLP0,RPS11,RPS15,RPS24,RPS8 | | | | | | | | |
| LXR/RXR Activation | | 4.54E+00 | | ALB,CLU,FDFT1,LPL,LYZ,PLTP,SAA1,SERPINA1,TF,VTN | | | | | | | | |
| FXR/RXR Activation | | 4.45E+00 | | ALB,CLU,CYP8B1,HNF1A,LPL,PLTP,SAA1,SERPINA1,TF,VTN | | | | | | | | |
| Acute Phase Response Signaling | | 3.70E+00 | | ALB,C1QA,C1QC,CEBPB,FTL,HMOX2,HNF1A,SAA1,SAA2-SAA4,SERPINA1,TF | | | | | | | | |
| Adipogenesis pathway | | 2.82E+00 | | CEBPB,DDIT3,ERCC2,FABP4,LPIN1,LPL,RBBP4,TXNIP | | | | | | | | |
| **Top significant upstream regulators** | | | | | | | | | | | | |
| **Upstream Regulator** | **Molecule Type** | | | | **Predicted Activation State** | | | **Activation z-score** | **p-value of overlap** | | | |
| ACOX1 | enzyme | | | | Inhibited | | | -3.105 | 3.70E-16 | | | |
| MLXIPL | transcription regulator | | | | Activated | | | 3.137 | 6.99E-12 | | | |
| ciprofibrate | chemical drug | | | |  | | | 1.958 | 5.00E-11 | | | |
| LARP1 | translation regulator | | | | Inhibited | | | -3.742 | 1.02E-10 | | | |
| MYCN | transcription regulator | | | | Activated | | | 2.324 | 1.09E-10 | | | |
| **Top significant diseases and disorders** | | | | | | | | | | | | |
| **Categories** | | | | | | | **Diseases.or.Functions.Annotation** | | | | | **p-value** |
| Cancer,Organismal Injury and Abnormalities | | | | | | | Non-melanoma solid tumor | | | | | 5.74E-12 |
| Cancer,Organismal Injury and Abnormalities | | | | | | | Epithelial neoplasm | | | | | 6.61E-11 |
| Cancer,Organismal Injury and Abnormalities | | | | | | | Non-hematological solid tumor | | | | | 9.74E-11 |
| Cancer,Organismal Injury and Abnormalities | | | | | | | Nonhematologic malignant neoplasm | | | | | 1.13E-10 |
| Cancer,Organismal Injury and Abnormalities | | | | | | | Carcinoma | | | | | 1.3E-10 |
| Cancer,Organismal Injury and Abnormalities | | | | | | | Malignant solid tumor | | | | | 1.58E-10 |
| Cancer,Organismal Injury and Abnormalities | | | | | | | Solid tumor | | | | | 1.97E-10 |
| Cancer,Organismal Injury and Abnormalities | | | | | | | Extracranial solid tumor | | | | | 1.9E-09 |
| Cellular Compromise,Inflammatory Response | | | | | | | Degranulation of phagocytes | | | | | 5.92E-08 |
| Cellular Compromise,Inflammatory Response | | | | | | | Degranulation of cells | | | | | 6.7E-08 |
| Developmental Disorder,Hereditary Disorder,Metabolic Disease,Organismal Injury and Abnormalities | | | | | | | Familial amyloidosis | | | | | 7.84E-08 |
| Cellular Compromise,Inflammatory Response | | | | | | | Degranulation of myeloid cells | | | | | 8.06E-08 |
| Organismal Injury and Abnormalities | | | | | | | Benign lesion | | | | | 9.73E-08 |
| Cancer,Organismal Injury and Abnormalities | | | | | | | Intraabdominal organ tumor | | | | | 1.32E-07 |
| Cancer,Organismal Injury and Abnormalities | | | | | | | Head and neck cancer | | | | | 1.44E-07 |
| Cancer,Organismal Injury and Abnormalities | | | | | | | Cancer of secretory structure | | | | | 1.67E-07 |
| Developmental Disorder,Neurological Disease | | | | | | | Congenital neurological disorder | | | | | 2.14E-07 |
| Cancer,Gastrointestinal Disease,Organismal Injury and Abnormalities | | | | | | | Digestive organ tumor | | | | | 2.64E-07 |
| Cellular Movement,Hematological System Development and Function,Immune Cell Trafficking,Inflammatory Response | | | | | | | Chemotaxis of neutrophils | | | | | 2.79E-07 |
| Cancer,Organismal Injury and Abnormalities | | | | | | | Head and neck tumor | | | | | 2.93E-07 |
| Cancer,Organismal Injury and Abnormalities | | | | | | | Abdominal neoplasm | | | | | 5.34E-07 |
| Cancer,Endocrine System Disorders,Organismal Injury and Abnormalities | | | | | | | Endocrine carcinoma | | | | | 6.83E-07 |
| Organismal Development,Organismal Injury and Abnormalities | | | | | | | Abnormal morphology of abdomen | | | | | 7.02E-07 |
| Metabolic Disease,Organismal Injury and Abnormalities | | | | | | | Amyloidosis | | | | | 7.21E-07 |
| Cancer,Endocrine System Disorders,Organismal Injury and Abnormalities | | | | | | | Endocrine gland tumor | | | | | 8.23E-07 |
| Cancer,Organismal Injury and Abnormalities | | | | | | | Adenocarcinoma | | | | | 9.27E-07 |
| Dermatological Diseases and Conditions,Immunological Disease,Inflammatory Disease,Inflammatory Response,Organismal Injury and Abnormalities | | | | | | | Atopic dermatitis | | | | | 0.00000121 |
| Gastrointestinal Disease,Inflammatory Disease,Inflammatory Response,Organismal Injury and Abnormalities | | | | | | | Chronic colitis | | | | | 0.00000128 |
| Cancer,Organismal Injury and Abnormalities | | | | | | | Neck neoplasm | | | | | 0.00000162 |
| Cancer,Cell Death and Survival,Organismal Injury and Abnormalities,Tumor Morphology | | | | | | | Cell death of osteosarcoma cells | | | | | 0.00000198 |
| Cancer,Endocrine System Disorders,Organismal Injury and Abnormalities | | | | | | | Nonpituitary endocrine tumor | | | | | 0.00000204 |
| Cellular Compromise,Inflammatory Response | | | | | | | Degranulation of neutrophils | | | | | 0.00000211 |
| Organismal Injury and Abnormalities,Renal and Urological Disease | | | | | | | Proximal tubular toxicity | | | | | 0.00000238 |
| Cellular Movement,Hematological System Development and Function,Immune Cell Trafficking,Inflammatory Response | | | | | | | Cell movement of neutrophils | | | | | 0.00000243 |
| Cancer,Organismal Injury and Abnormalities | | | | | | | Head and neck carcinoma | | | | | 0.00000257 |
| Cancer,Endocrine System Disorders,Organismal Injury and Abnormalities | | | | | | | Thyroid carcinoma | | | | | 0.00000263 |
| Cancer,Endocrine System Disorders,Organismal Injury and Abnormalities | | | | | | | Thyroid gland tumor | | | | | 0.0000027 |
| Cancer,Organismal Injury and Abnormalities | | | | | | | Benign solid tumor | | | | | 0.00000297 |
| Digestive System Development and Function,Gastrointestinal Disease,Hepatic System Development and Function,Hepatic System Disease,Organ Morphology,Organismal Development,Organismal Injury and Abnormalities | | | | | | | Abnormal morphology of liver | | | | | 0.00000303 |
| Cancer,Organismal Injury and Abnormalities | | | | | | | Abdominal adenocarcinoma | | | | | 0.00000369 |
| Hereditary Disorder,Neurological Disease,Organismal Injury and Abnormalities | | | | | | | Autosomal recessive neurological disorder | | | | | 0.00000373 |
| Inflammatory Response | | | | | | | Inflammatory response | | | | | 0.00000404 |
| Cancer,Organismal Injury and Abnormalities | | | | | | | Abdominal carcinoma | | | | | 0.00000456 |
| Metabolic Disease,Organismal Injury and Abnormalities | | | | | | | Glucose metabolism disorder | | | | | 0.0000049 |
| Cellular Movement,Hematological System Development and Function,Immune Cell Trafficking,Inflammatory Response | | | | | | | Chemotaxis of phagocytes | | | | | 0.00000518 |
| Cancer,Cell Death and Survival,Organismal Injury and Abnormalities,Tumor Morphology | | | | | | | Necrosis of malignant tumor | | | | | 0.00000533 |
| Inflammatory Response,Organismal Injury and Abnormalities | | | | | | | Inflammation of organ | | | | | 0.0000065 |
| Cancer,Cell Death and Survival,Organismal Injury and Abnormalities,Tumor Morphology | | | | | | | Necrosis of tumor | | | | | 0.00000799 |
| Cancer,Hematological Disease,Immunological Disease,Organismal Injury and Abnormalities | | | | | | | Lymphoma | | | | | 0.00000896 |
| Cell-To-Cell Signaling and Interaction,Hematological System Development and Function,Immune Cell Trafficking,Inflammatory Response | | | | | | | Binding of neutrophils | | | | | 0.00000917 |
| Connective Tissue Disorders,Inflammatory Disease,Organismal Injury and Abnormalities,Skeletal and Muscular Disorders | | | | | | | Rheumatic Disease | | | | | 0.0000105 |
| Organismal Injury and Abnormalities,Tissue Morphology | | | | | | | Morphology of lesion | | | | | 0.0000109 |
| Cancer,Gastrointestinal Disease,Organismal Injury and Abnormalities | | | | | | | Digestive system cancer | | | | | 0.0000114 |
| Neurological Disease,Organismal Injury and Abnormalities,Psychological Disorders | | | | | | | Tauopathy | | | | | 0.0000116 |
| Cancer,Organismal Injury and Abnormalities | | | | | | | Papillary carcinoma | | | | | 0.0000129 |
| Cancer,Cell Death and Survival,Organismal Injury and Abnormalities,Tumor Morphology | | | | | | | Cell death of cancer cells | | | | | 0.0000156 |
| Cancer,Organismal Injury and Abnormalities | | | | | | | Abdominal cancer | | | | | 0.0000161 |
| Organismal Injury and Abnormalities,Tissue Morphology | | | | | | | Size of lesion | | | | | 0.0000178 |
| Cancer,Organismal Injury and Abnormalities | | | | | | | Cancer of cells | | | | | 0.0000179 |
| Cancer,Organismal Injury and Abnormalities | | | | | | | Formation of solid tumor | | | | | 0.000019 |
| Developmental Disorder,Hereditary Disorder,Metabolic Disease,Neurological Disease,Organismal Injury and Abnormalities | | | | | | | Familial amyloidotic polyneuropathy | | | | | 0.0000193 |
| Cancer,Organismal Injury and Abnormalities | | | | | | | Neoplasia of cells | | | | | 0.0000216 |
| Cancer,Organismal Injury and Abnormalities,Renal and Urological Disease | | | | | | | Urinary tract cancer | | | | | 0.0000231 |
| Cancer,Organismal Injury and Abnormalities | | | | | | | Metastatic carcinoma | | | | | 0.0000259 |
| Cancer,Gastrointestinal Disease,Organismal Injury and Abnormalities | | | | | | | Gastrointestinal adenocarcinoma | | | | | 0.0000272 |
| Cancer,Organismal Injury and Abnormalities,Renal and Urological Disease | | | | | | | Urinary tract tumor | | | | | 0.0000282 |
| Connective Tissue Disorders,Inflammatory Disease,Inflammatory Response,Organismal Injury and Abnormalities,Skeletal and Muscular Disorders | | | | | | | Inflammation of joint | | | | | 0.0000287 |
| Cancer,Organismal Injury and Abnormalities | | | | | | | Incidence of tumor | | | | | 0.0000288 |
| Hematological System Development and Function,Immunological Disease,Lymphoid Tissue Structure and Development,Organismal Injury and Abnormalities,Tissue Morphology | | | | | | | Abnormal morphology of B-cell follicle | | | | | 0.0000296 |
| Organismal Injury and Abnormalities,Renal and Urological Disease | | | | | | | Urination disorder | | | | | 0.0000309 |
| Cancer,Organismal Injury and Abnormalities | | | | | | | Papillary adenocarcinoma | | | | | 0.0000313 |
| Cancer,Gastrointestinal Disease,Organismal Injury and Abnormalities | | | | | | | Gastrointestinal carcinoma | | | | | 0.0000325 |
| Cancer,Gastrointestinal Disease,Organismal Injury and Abnormalities | | | | | | | Gastrointestinal tumor | | | | | 0.0000345 |
| Cancer,Organismal Injury and Abnormalities | | | | | | | Frequency of tumor | | | | | 0.0000357 |
| Metabolic Disease,Organismal Injury and Abnormalities | | | | | | | Disorder of lipid metabolism | | | | | 0.000038 |
| Cancer,Cell Death and Survival,Organismal Injury and Abnormalities,Tumor Morphology | | | | | | | Cell death of tumor cells | | | | | 0.0000421 |
| Cancer,Organismal Injury and Abnormalities | | | | | | | Lymphoreticular neoplasm | | | | | 0.0000445 |
| Neurological Disease,Organismal Injury and Abnormalities | | | | | | | Progressive encephalopathy | | | | | 0.0000471 |
| Cellular Compromise,Inflammatory Response | | | | | | | Degranulation of blood platelets | | | | | 0.0000498 |
| Inflammatory Response | | | | | | | Inflammation of secretory structure | | | | | 0.0000502 |
| Endocrine System Disorders,Metabolic Disease,Organismal Injury and Abnormalities | | | | | | | Insulin resistance | | | | | 0.0000518 |
| Cancer,Organismal Injury and Abnormalities | | | | | | | Tumorigenesis of epithelial neoplasm | | | | | 0.0000529 |
| Endocrine System Disorders,Gastrointestinal Disease,Inflammatory Disease,Inflammatory Response,Organismal Injury and Abnormalities | | | | | | | Inflammation of pancreas | | | | | 0.0000529 |
| Cancer,Organismal Injury and Abnormalities | | | | | | | Mucoepidermoid carcinoma | | | | | 0.0000579 |
| Cardiovascular Disease,Hematological Disease,Metabolic Disease,Organismal Injury and Abnormalities | | | | | | | Hyperlipidemia | | | | | 0.0000586 |
| Cancer,Gastrointestinal Disease,Organismal Injury and Abnormalities | | | | | | | Gastrointestinal tract cancer | | | | | 0.0000602 |
| Cancer,Organismal Injury and Abnormalities | | | | | | | Growth of tumor | | | | | 0.0000603 |
| Hematological Disease,Metabolic Disease,Organismal Injury and Abnormalities | | | | | | | Dyslipidemia | | | | | 0.0000605 |
| Gastrointestinal Disease,Inflammatory Disease,Inflammatory Response,Organismal Injury and Abnormalities | | | | | | | Experimental colitis | | | | | 0.0000605 |
| Cancer,Organismal Injury and Abnormalities | | | | | | | Advanced stage carcinoma | | | | | 0.0000626 |
| Cancer,Organismal Injury and Abnormalities | | | | | | | Development of malignant tumor | | | | | 0.0000772 |
| Cellular Movement,Hematological System Development and Function,Immune Cell Trafficking,Inflammatory Response | | | | | | | Cell movement of phagocytes | | | | | 0.0000775 |
| Cell-To-Cell Signaling and Interaction,Hematological System Development and Function,Immune Cell Trafficking,Inflammatory Response | | | | | | | Adhesion of neutrophils | | | | | 0.0000794 |
| Dermatological Diseases and Conditions,Organismal Injury and Abnormalities | | | | | | | Psoriasis | | | | | 0.00008 |
| Inflammatory Response | | | | | | | Inflammation of absolute anatomical region | | | | | 0.0000813 |
| Cancer,Organismal Injury and Abnormalities | | | | | | | Development of carcinoma | | | | | 0.0000832 |
| Inflammatory Response | | | | | | | Inflammation of body cavity | | | | | 0.0000866 |
| Cell-To-Cell Signaling and Interaction,Hematological System Development and Function,Immune Cell Trafficking,Inflammatory Response | | | | | | | Activation of leukocytes | | | | | 0.0000877 |
| Cancer,Organismal Injury and Abnormalities | | | | | | | Invasive tumor | | | | | 0.000088 |
| Cancer,Organismal Injury and Abnormalities | | | | | | | Pelvic tumor | | | | | 0.000088 |
| Hereditary Disorder,Neurological Disease,Organismal Injury and Abnormalities | | | | | | | Peroxisomal acyl CoA oxidase deficiency | | | | | 0.0001 |
| Gastrointestinal Disease,Hepatic System Disease,Organismal Injury and Abnormalities | | | | | | | Liver lesion | | | | | 0.000106 |
| Cellular Movement,Hematological System Development and Function,Immune Cell Trafficking,Inflammatory Response | | | | | | | Migration of neutrophils | | | | | 0.000106 |
| Cancer,Organismal Injury and Abnormalities | | | | | | | Malignant genitourinary solid tumor | | | | | 0.000107 |
| Developmental Disorder,Hereditary Disorder,Metabolic Disease,Neurological Disease,Organismal Injury and Abnormalities,Psychological Disorders | | | | | | | Familial Alzheimer disease | | | | | 0.000111 |
| Cancer,Gastrointestinal Disease,Organismal Injury and Abnormalities | | | | | | | Large intestine carcinoma | | | | | 0.000115 |
| Cancer,Organismal Injury and Abnormalities | | | | | | | Pelvic cancer | | | | | 0.000121 |
| Organismal Injury and Abnormalities | | | | | | | Fibrosis | | | | | 0.000131 |
| Gastrointestinal Disease,Hepatic System Disease,Metabolic Disease,Organismal Injury and Abnormalities | | | | | | | Hepatic steatosis | | | | | 0.000131 |
| Nervous System Development and Function,Neurological Disease,Organ Morphology,Organismal Development,Organismal Injury and Abnormalities,Psychological Disorders | | | | | | | Abnormal morphology of substantia nigra | | | | | 0.000135 |
| Dermatological Diseases and Conditions,Organismal Injury and Abnormalities | | | | | | | Benign skin lesion | | | | | 0.000135 |
| Cancer,Organismal Injury and Abnormalities | | | | | | | Lymphatic system tumor | | | | | 0.000136 |
| Cancer,Organismal Injury and Abnormalities,Respiratory Disease | | | | | | | Stage I metastatic non-small cell lung carcinoma | | | | | 0.000143 |
| Hematological System Development and Function,Immunological Disease,Lymphoid Tissue Structure and Development,Organ Morphology,Organismal Injury and Abnormalities,Tissue Morphology | | | | | | | Abnormal morphology of lymph node | | | | | 0.000144 |
| Neurological Disease,Organismal Injury and Abnormalities,Psychological Disorders | | | | | | | Alzheimer disease or frontotemporal dementia | | | | | 0.000145 |
| Cancer,Organismal Injury and Abnormalities | | | | | | | Genitourinary tumor | | | | | 0.000161 |
| Organismal Injury and Abnormalities,Renal and Urological Disease,Renal and Urological System Development and Function | | | | | | | Abnormal morphology of urinary system | | | | | 0.000165 |
| Cancer,Organismal Injury and Abnormalities | | | | | | | Development of benign tumor | | | | | 0.000204 |
| Cancer,Gastrointestinal Disease,Organismal Injury and Abnormalities | | | | | | | Large intestine neoplasm | | | | | 0.000206 |
| Connective Tissue Disorders,Immunological Disease,Inflammatory Disease,Inflammatory Response,Organismal Injury and Abnormalities,Skeletal and Muscular Disorders | | | | | | | Rheumatoid arthritis | | | | | 0.000215 |
| Dermatological Diseases and Conditions,Organismal Injury and Abnormalities | | | | | | | Abnormality of skin morphology | | | | | 0.000235 |
| Metabolic Disease,Neurological Disease,Organismal Injury and Abnormalities,Psychological Disorders | | | | | | | Alzheimer disease | | | | | 0.000245 |
| Cancer,Organismal Injury and Abnormalities,Tissue Morphology,Tumor Morphology | | | | | | | Morphology of tumor | | | | | 0.000246 |
| Developmental Disorder,Hereditary Disorder,Immunological Disease,Neurological Disease,Organismal Injury and Abnormalities,Skeletal and Muscular Disorders | | | | | | | Congenital myasthenic syndrome type 22 | | | | | 0.000254 |
| Endocrine System Disorders,Gastrointestinal Disease,Metabolic Disease,Organismal Injury and Abnormalities | | | | | | | Diabetes mellitus | | | | | 0.000256 |
| Dermatological Diseases and Conditions,Inflammatory Disease,Inflammatory Response,Organismal Injury and Abnormalities | | | | | | | Dermatitis | | | | | 0.000262 |
| Gastrointestinal Disease,Inflammatory Disease,Inflammatory Response,Organismal Injury and Abnormalities | | | | | | | Dextran sodium sulfate-induced colitis | | | | | 0.000265 |
| Hematological System Development and Function,Inflammatory Response,Tissue Morphology | | | | | | | Quantity of phagocytes | | | | | 0.000269 |
| Cancer,Hematological Disease,Immunological Disease,Organismal Injury and Abnormalities | | | | | | | Non-Hodgkin lymphoma | | | | | 0.000288 |
| Cancer,Hematological Disease,Immunological Disease,Organismal Injury and Abnormalities | | | | | | | Lymphocytic cancer | | | | | 0.0003 |
| Cancer,Organismal Injury and Abnormalities | | | | | | | Adenoma | | | | | 0.00031 |
| Cancer,Hematological Disease,Organismal Injury and Abnormalities | | | | | | | Lymphocytic neoplasm | | | | | 0.000313 |
| Cancer,Gastrointestinal Disease,Organismal Injury and Abnormalities | | | | | | | Large intestine adenocarcinoma | | | | | 0.000316 |
| Hereditary Disorder,Neurological Disease,Organismal Injury and Abnormalities | | | | | | | Familial encephalopathy | | | | | 0.000334 |
| Cancer,Gastrointestinal Disease,Organismal Injury and Abnormalities | | | | | | | Malignant neoplasm of large intestine | | | | | 0.000337 |
| Neurological Disease,Organismal Injury and Abnormalities,Psychological Disorders | | | | | | | Dementia | | | | | 0.000338 |
| Cancer,Organismal Injury and Abnormalities | | | | | | | Head and neck adenocarcinoma | | | | | 0.000339 |
| Cell-To-Cell Signaling and Interaction,Hematological System Development and Function,Inflammatory Response | | | | | | | Binding of professional phagocytic cells | | | | | 0.000364 |
| Cancer,Organismal Injury and Abnormalities | | | | | | | Metastatic solid tumor | | | | | 0.000367 |
| Cancer,Hematological Disease,Organismal Injury and Abnormalities | | | | | | | Hematologic cancer of cells | | | | | 0.000373 |
| Cancer,Neurological Disease,Organismal Injury and Abnormalities | | | | | | | Brain oligodendroglioma | | | | | 0.000377 |
| Hematological System Development and Function,Immunological Disease,Lymphoid Tissue Structure and Development,Organ Morphology,Organismal Injury and Abnormalities,Tissue Morphology | | | | | | | Abnormal morphology of lymphoid organ | | | | | 0.000405 |
| Dermatological Diseases and Conditions,Organismal Injury and Abnormalities | | | | | | | Blister | | | | | 0.00042 |
| Cancer,Organismal Injury and Abnormalities,Renal and Urological Disease | | | | | | | Renal tumor | | | | | 0.000443 |
| Cancer,Organismal Injury and Abnormalities,Respiratory Disease | | | | | | | Advanced non-small cell lung carcinoma | | | | | 0.000443 |
| Developmental Disorder,Hereditary Disorder,Neurological Disease,Organismal Injury and Abnormalities | | | | | | | Smith-Magenis syndrome-like disorder | | | | | 0.000446 |
| Hematological System Development and Function,Immunological Disease,Lymphoid Tissue Structure and Development,Organ Morphology,Organismal Development,Organismal Injury and Abnormalities,Tissue Morphology | | | | | | | Abnormal morphology of periarteriolar lymphoid sheath | | | | | 0.000446 |
| Cancer,Endocrine System Disorders,Gastrointestinal Disease,Organismal Injury and Abnormalities | | | | | | | Growth of pancreatic endocrine tumor | | | | | 0.000446 |
| Cancer,Organismal Injury and Abnormalities | | | | | | | Squamous-cell carcinoma | | | | | 0.000449 |
| Cancer,Neurological Disease,Organismal Injury and Abnormalities | | | | | | | Grade 3 malignant glioma | | | | | 0.000462 |
| Cellular Movement,Hematological System Development and Function,Immune Cell Trafficking,Inflammatory Response | | | | | | | Migration of phagocytes | | | | | 0.000466 |
| Cancer,Hematological Disease,Organismal Injury and Abnormalities | | | | | | | Myeloid or lymphoid neoplasm | | | | | 0.000469 |
| Cancer,Organismal Injury and Abnormalities,Renal and Urological Disease | | | | | | | Renal cancer | | | | | 0.000472 |
| Cancer,Neurological Disease,Organismal Injury and Abnormalities | | | | | | | Grade 2-3 glioma | | | | | 0.000499 |
| Organismal Injury and Abnormalities,Tissue Morphology | | | | | | | Abnormal morphology of epithelial tissue | | | | | 0.000499 |
| Developmental Disorder,Hereditary Disorder,Organismal Injury and Abnormalities,Skeletal and Muscular Disorders | | | | | | | Duchenne muscular dystrophy | | | | | 0.0005 |
| Connective Tissue Disorders,Organismal Injury and Abnormalities,Skeletal and Muscular Disorders | | | | | | | Non-traumatic arthropathy | | | | | 0.000507 |
| Cancer,Neurological Disease,Organismal Injury and Abnormalities | | | | | | | Oligodendroglioma | | | | | 0.000509 |
| Cancer,Hematological Disease,Organismal Injury and Abnormalities | | | | | | | Hematologic cancer | | | | | 0.000512 |
| Connective Tissue Disorders,Immunological Disease,Inflammatory Disease,Organismal Injury and Abnormalities,Skeletal and Muscular Disorders | | | | | | | Lupus erythematosus | | | | | 0.00052 |
| Endocrine System Disorders,Gastrointestinal Disease,Metabolic Disease,Organismal Injury and Abnormalities | | | | | | | Non-insulin-dependent diabetes mellitus | | | | | 0.000521 |
| Cancer,Organismal Injury and Abnormalities | | | | | | | Advanced malignant solid tumor | | | | | 0.000524 |
| Organismal Injury and Abnormalities,Reproductive System Disease | | | | | | | Benign pelvic disease | | | | | 0.00053 |
| Neurological Disease,Organismal Injury and Abnormalities,Psychological Disorders | | | | | | | Disorder of basal ganglia | | | | | 0.000532 |
| Cancer,Hematological Disease,Immunological Disease,Organismal Injury and Abnormalities | | | | | | | Neoplasia of leukocytes | | | | | 0.000539 |
| Cell-To-Cell Signaling and Interaction,Hematological System Development and Function,Immune Cell Trafficking,Inflammatory Response | | | | | | | Activation of lymphocytes | | | | | 0.000541 |
| Hereditary Disorder,Immunological Disease,Organismal Injury and Abnormalities | | | | | | | Autosomal recessive immunological disorder | | | | | 0.000553 |
| Cancer,Hematological Disease,Immunological Disease,Organismal Injury and Abnormalities | | | | | | | Tumorigenesis of lymphocytes | | | | | 0.000556 |
| Cancer,Gastrointestinal Disease,Organismal Injury and Abnormalities | | | | | | | Non-colon gastrointestinal cancer | | | | | 0.000557 |
| Cancer,Hematological Disease,Immunological Disease,Organismal Injury and Abnormalities | | | | | | | B-cell lymphoma | | | | | 0.00057 |
| Cancer,Organismal Injury and Abnormalities,Respiratory Disease | | | | | | | Lung tumor | | | | | 0.000597 |
| Cellular Movement,Hematological System Development and Function,Immune Cell Trafficking,Inflammatory Response | | | | | | | Cell movement of macrophages | | | | | 0.000606 |
| Cancer,Cellular Development,Cellular Growth and Proliferation,Organismal Injury and Abnormalities,Tumor Morphology | | | | | | | Proliferation of cancer cells | | | | | 0.000609 |
| Developmental Disorder,Hereditary Disorder,Organismal Injury and Abnormalities,Skeletal and Muscular Disorders | | | | | | | Dystrophy of muscle | | | | | 0.00062 |
| Infectious Diseases,Neurological Disease,Organismal Injury and Abnormalities,Psychological Disorders | | | | | | | Prion disease | | | | | 0.000621 |
| Cell Death and Survival,Organismal Injury and Abnormalities,Renal and Urological Disease | | | | | | | Apoptosis of kidney cells | | | | | 0.000624 |
| Cell-To-Cell Signaling and Interaction,Cellular Function and Maintenance,Hematological System Development and Function,Inflammatory Response | | | | | | | Phagocytosis by macrophages | | | | | 0.000634 |
| Developmental Disorder | | | | | | | Growth failure or short stature | | | | | 0.000669 |
| Cancer,Gastrointestinal Disease,Organismal Injury and Abnormalities | | | | | | | Upper gastrointestinal tract tumor | | | | | 0.000675 |
| Cancer,Organismal Injury and Abnormalities,Respiratory Disease | | | | | | | Advanced lung cancer | | | | | 0.000678 |
| Cancer,Organismal Injury and Abnormalities | | | | | | | Metastasis | | | | | 0.000703 |
| Cancer,Organismal Injury and Abnormalities | | | | | | | Breast or pancreatic cancer | | | | | 0.00071 |
| Cancer,Organismal Injury and Abnormalities,Respiratory Disease | | | | | | | Lung carcinoma | | | | | 0.00073 |
| Hematological System Development and Function,Immune Cell Trafficking,Inflammatory Response,Tissue Development | | | | | | | Accumulation of neutrophils | | | | | 0.000731 |
| Hematological System Development and Function,Immunological Disease,Lymphoid Tissue Structure and Development,Organ Morphology,Organismal Development,Organismal Injury and Abnormalities,Tissue Morphology | | | | | | | Abnormal morphology of spleen | | | | | 0.000744 |
| Organismal Injury and Abnormalities,Renal and Urological Disease | | | | | | | Renal lesion | | | | | 0.000749 |
| Inflammatory Disease,Inflammatory Response,Organismal Injury and Abnormalities,Renal and Urological Disease | | | | | | | Glomerulonephritis | | | | | 0.000749 |
| Metabolic Disease,Organismal Injury and Abnormalities | | | | | | | Hypovolemia | | | | | 0.000755 |
| Developmental Disorder,Embryonic Development,Organismal Development,Tissue Morphology | | | | | | | Lack of first branchial arch | | | | | 0.000755 |
| Developmental Disorder,Hereditary Disorder,Immunological Disease,Organismal Injury and Abnormalities | | | | | | | Complement component C1q deficiency | | | | | 0.000755 |
| Cancer,Organismal Injury and Abnormalities,Reproductive System Disease | | | | | | | Breast cancer | | | | | 0.000795 |
| Cell-To-Cell Signaling and Interaction,Cellular Function and Maintenance,Inflammatory Response | | | | | | | Phagocytosis of phagocytes | | | | | 0.000806 |
| Cancer,Organismal Injury and Abnormalities,Reproductive System Disease | | | | | | | Breast or ovarian cancer | | | | | 0.000814 |
| Cancer,Hematological Disease,Organismal Injury and Abnormalities | | | | | | | Neoplasia of blood cells | | | | | 0.000816 |
| Cancer,Organismal Injury and Abnormalities | | | | | | | Pelvic carcinoma | | | | | 0.000817 |
| Cancer,Organismal Injury and Abnormalities,Respiratory Disease | | | | | | | Lung cancer | | | | | 0.000825 |
| Cancer,Gastrointestinal Disease,Organismal Injury and Abnormalities | | | | | | | Upper gastrointestinal tract cancer | | | | | 0.000833 |
| Organismal Injury and Abnormalities,Reproductive System Disease | | | | | | | Benign uterine disease | | | | | 0.000849 |
| Cancer,Organismal Injury and Abnormalities | | | | | | | Advanced extracranial solid tumor | | | | | 0.00086 |
| Organismal Injury and Abnormalities | | | | | | | Damage of genitourinary system | | | | | 0.000873 |
| Organismal Injury and Abnormalities,Respiratory Disease | | | | | | | Chronic obstructive pulmonary disease | | | | | 0.000904 |
| Cell-To-Cell Signaling and Interaction,Hematological System Development and Function,Immune Cell Trafficking,Inflammatory Response | | | | | | | Adhesion of phagocytes | | | | | 0.000925 |
| Cancer,Organismal Injury and Abnormalities | | | | | | | Melanoma | | | | | 0.000949 |
| Endocrine System Disorders,Gastrointestinal Disease,Inflammatory Disease,Inflammatory Response,Organismal Injury and Abnormalities | | | | | | | Insulitis | | | | | 0.000985 |
| Cancer,Organismal Injury and Abnormalities,Tissue Morphology,Tumor Morphology | | | | | | | Size of tumor | | | | | 0.000999 |
| Cancer,Organismal Injury and Abnormalities | | | | | | | Benign connective or soft tissue neoplasm | | | | | 0.001 |
| Hematological Disease,Immunological Disease,Organismal Injury and Abnormalities | | | | | | | Leukocytosis | | | | | 0.00102 |
| Gastrointestinal Disease,Inflammatory Response | | | | | | | Inflammation of gastrointestinal tract | | | | | 0.00104 |
| Cancer,Gastrointestinal Disease,Organismal Injury and Abnormalities | | | | | | | Development of digestive organ tumor | | | | | 0.00104 |
| Digestive System Development and Function,Gastrointestinal Disease,Hepatic System Development and Function,Hepatic System Disease,Organ Morphology,Organismal Development,Organismal Injury and Abnormalities | | | | | | | Hepatomegaly | | | | | 0.00106 |
| Cancer,Organismal Injury and Abnormalities | | | | | | | Growth of malignant tumor | | | | | 0.00108 |
| Dermatological Diseases and Conditions,Organismal Injury and Abnormalities | | | | | | | Exanthem of skin | | | | | 0.00109 |
| Cancer,Organismal Injury and Abnormalities | | | | | | | Malignant neoplasm of retroperitoneum | | | | | 0.00109 |
| Neurological Disease,Organismal Injury and Abnormalities | | | | | | | Demyelination of brain | | | | | 0.00111 |
| Developmental Disorder | | | | | | | Growth Failure | | | | | 0.00116 |
| Cellular Movement,Hematological System Development and Function,Immune Cell Trafficking,Inflammatory Response | | | | | | | Transmigration of monocytes | | | | | 0.00119 |
| Cell Death and Survival,Organismal Injury and Abnormalities,Renal and Urological Disease | | | | | | | Apoptosis of tubular cells | | | | | 0.00119 |
| Cancer,Organismal Injury and Abnormalities,Renal and Urological Disease | | | | | | | Bladder cancer | | | | | 0.00119 |
| Cardiovascular Disease,Organismal Injury and Abnormalities,Tissue Morphology | | | | | | | Size of vascular lesion | | | | | 0.00119 |
| Hematological System Development and Function,Inflammatory Response,Tissue Morphology | | | | | | | Quantity of monocytes | | | | | 0.00122 |
| Cellular Movement,Hematological System Development and Function,Immune Cell Trafficking,Inflammatory Response | | | | | | | Chemotaxis of macrophages | | | | | 0.00124 |
| Cancer,Endocrine System Disorders,Organismal Injury and Abnormalities | | | | | | | Thyroid gland nonmedullary carcinoma | | | | | 0.00124 |
| Connective Tissue Development and Function,Connective Tissue Disorders,Organ Morphology,Organismal Injury and Abnormalities,Skeletal and Muscular Disorders,Skeletal and Muscular System Development and Function,Tissue Development | | | | | | | Abnormal morphology of metaphysis | | | | | 0.00126 |
| Cancer,Organismal Injury and Abnormalities,Respiratory Disease | | | | | | | Metastatic non-small cell lung carcinoma | | | | | 0.00127 |
| Connective Tissue Development and Function,Connective Tissue Disorders,Organismal Injury and Abnormalities,Skeletal and Muscular Disorders,Skeletal and Muscular System Development and Function,Tissue Development | | | | | | | Abnormal morphology of diaphysis | | | | | 0.00129 |
| Hematological System Development and Function,Inflammatory Response,Tissue Morphology | | | | | | | Quantity of neutrophils | | | | | 0.0013 |
| Cancer,Organismal Injury and Abnormalities | | | | | | | Advanced malignant tumor | | | | | 0.00133 |
| Cancer,Immunological Disease,Organismal Injury and Abnormalities | | | | | | | Hyperplasia of lymphoid organ | | | | | 0.00134 |
| Cell-To-Cell Signaling and Interaction,Inflammatory Response | | | | | | | Immune response of antigen presenting cells | | | | | 0.00134 |
| Cancer,Cellular Development,Cellular Growth and Proliferation,Organismal Injury and Abnormalities,Tumor Morphology | | | | | | | Proliferation of tumor cells | | | | | 0.00135 |
| Hematological System Development and Function,Immune Cell Trafficking,Inflammatory Response,Tissue Development | | | | | | | Accumulation of granulocytes | | | | | 0.00136 |
| Inflammatory Disease,Inflammatory Response,Organismal Injury and Abnormalities,Renal and Urological Disease | | | | | | | Nephritis | | | | | 0.00138 |
| Ophthalmic Disease,Organismal Injury and Abnormalities | | | | | | | Retinal degeneration | | | | | 0.00139 |
| Cancer,Organismal Injury and Abnormalities | | | | | | | Blue round small cell tumor | | | | | 0.00139 |
| Cancer,Organismal Injury and Abnormalities,Reproductive System Disease,Skeletal and Muscular Disorders | | | | | | | Uterine leiomyoma | | | | | 0.00141 |
| Dermatological Diseases and Conditions,Immunological Disease,Inflammatory Disease,Organismal Injury and Abnormalities | | | | | | | Lichen planus | | | | | 0.00141 |
| Organ Morphology,Organismal Development,Organismal Injury and Abnormalities,Renal and Urological Disease,Renal and Urological System Development and Function | | | | | | | Abnormal morphology of urinary bladder | | | | | 0.00143 |
| Cancer,Organismal Injury and Abnormalities,Reproductive System Disease | | | | | | | Mammary tumor | | | | | 0.00144 |
| Cell Death and Survival,Organismal Injury and Abnormalities,Renal and Urological Disease | | | | | | | Necrosis of renal tubule | | | | | 0.00146 |
| Cancer,Endocrine System Disorders,Gastrointestinal Disease,Organismal Injury and Abnormalities,Tumor Morphology | | | | | | | Invasion of pancreatic endocrine tumor | | | | | 0.00149 |
| Dermatological Diseases and Conditions,Inflammatory Disease,Inflammatory Response,Organismal Injury and Abnormalities | | | | | | | Ulcerative dermatitis | | | | | 0.00149 |
| Dermatological Diseases and Conditions,Organismal Injury and Abnormalities | | | | | | | Plaque psoriasis | | | | | 0.0015 |
| Organismal Injury and Abnormalities,Renal and Urological Disease | | | | | | | Damage of kidney | | | | | 0.00152 |
| Cancer,Endocrine System Disorders,Organismal Injury and Abnormalities | | | | | | | Differentiated thyroid cancer | | | | | 0.00158 |
| Cancer,Organismal Injury and Abnormalities | | | | | | | Anogenital cancer | | | | | 0.0016 |
| Cellular Compromise,Hypersensitivity Response,Inflammatory Response | | | | | | | Degranulation of mast cells | | | | | 0.00163 |
| Developmental Disorder,Hereditary Disorder,Organismal Injury and Abnormalities,Skeletal and Muscular Disorders | | | | | | | Progressive muscular dystrophy | | | | | 0.00163 |
| Hereditary Disorder,Neurological Disease,Organismal Injury and Abnormalities,Psychological Disorders | | | | | | | Familial dementia | | | | | 0.00166 |
| Cancer,Organismal Injury and Abnormalities | | | | | | | Genitourinary carcinoma | | | | | 0.00167 |
| Cellular Compromise,Hypersensitivity Response,Inflammatory Response | | | | | | | Degranulation of bone marrow-derived mast cells | | | | | 0.00168 |
| Cellular Movement,Hematological System Development and Function,Immune Cell Trafficking,Inflammatory Response | | | | | | | Chemotaxis of antigen presenting cells | | | | | 0.00171 |
| Cancer,Gastrointestinal Disease,Organismal Injury and Abnormalities | | | | | | | Esophageal squamous cell carcinoma | | | | | 0.00173 |
| Gastrointestinal Disease,Inflammatory Disease,Inflammatory Response,Organismal Injury and Abnormalities | | | | | | | Enteritis | | | | | 0.00177 |
| Cancer,Organismal Injury and Abnormalities | | | | | | | Malignant solid organ tumor | | | | | 0.00184 |
| Cardiovascular Disease,Organismal Injury and Abnormalities | | | | | | | Abnormality of heart ventricle | | | | | 0.00199 |
| Connective Tissue Disorders,Immunological Disease,Inflammatory Disease,Organismal Injury and Abnormalities,Skeletal and Muscular Disorders | | | | | | | Systemic lupus erythematosus | | | | | 0.00202 |
| Cardiovascular Disease,Hereditary Disorder,Organismal Injury and Abnormalities | | | | | | | Familial cardiovascular disease | | | | | 0.00203 |
| Gastrointestinal Disease,Inflammatory Disease,Inflammatory Response,Organismal Injury and Abnormalities | | | | | | | Colitis | | | | | 0.00203 |
| Hematological Disease,Metabolic Disease,Nutritional Disease,Organismal Injury and Abnormalities | | | | | | | Iron overload | | | | | 0.00203 |
| Cardiovascular Disease,Organismal Injury and Abnormalities | | | | | | | Myocardial dysfunction | | | | | 0.00204 |
| Cancer,Organismal Injury and Abnormalities | | | | | | | Malignant neoplasm of aerodigestive tract | | | | | 0.00213 |
| Cancer,Gastrointestinal Disease,Organismal Injury and Abnormalities | | | | | | | Upper gastrointestinal carcinoma | | | | | 0.00215 |
| Cancer,Organismal Injury and Abnormalities | | | | | | | Multiple cancers | | | | | 0.00216 |
| Organismal Injury and Abnormalities,Renal and Urological Disease | | | | | | | Acute kidney injury | | | | | 0.00216 |
| Cancer,Organismal Injury and Abnormalities | | | | | | | Connective or soft tissue tumor | | | | | 0.00219 |
| Cell Death and Survival,Organismal Injury and Abnormalities | | | | | | | Necrosis of epithelial tissue | | | | | 0.00221 |
| **Top significant molecular and cellular functions** | | | | | | | | | | | | |
| Categories | | | | | | Diseases.or.Functions.Annotation | | | | | p-value | |
| RNA Damage and Repair | | | | | | Nonsense-mediated mRNA decay | | | | | 2.85E-09 | |
| Cellular Development,Cellular Growth and Proliferation | | | | | | Proliferation of blood cells | | | | | 8.09E-09 | |
| Protein Synthesis | | | | | | Metabolism of protein | | | | | 2.01E-08 | |
| Cellular Development,Cellular Growth and Proliferation,Hematological System Development and Function,Lymphoid Tissue Structure and Development | | | | | | Proliferation of immune cells | | | | | 3.5E-08 | |
| Cell Death and Survival | | | | | | Necrosis | | | | | 4.53E-08 | |
| Cellular Growth and Proliferation,Lymphoid Tissue Structure and Development | | | | | | Proliferation of lymphatic system cells | | | | | 4.59E-08 | |
| Protein Synthesis | | | | | | Synthesis of protein | | | | | 8.12E-08 | |
| Cellular Development,Cellular Growth and Proliferation,Hematological System Development and Function,Lymphoid Tissue Structure and Development | | | | | | Proliferation of lymphocytes | | | | | 8.61E-08 | |
| Protein Synthesis | | | | | | Initiation of translation of protein | | | | | 9.86E-08 | |
| Protein Synthesis | | | | | | Translation | | | | | 7.12E-07 | |
| Cellular Growth and Proliferation,Connective Tissue Development and Function,Tissue Development | | | | | | Proliferation of connective tissue cells | | | | | 7.19E-07 | |
| Cell Death and Survival | | | | | | Apoptosis | | | | | 0.00000112 | |
| Cancer,Cell Death and Survival,Organismal Injury and Abnormalities,Tumor Morphology | | | | | | Cell death of osteosarcoma cells | | | | | 0.00000198 | |
| Protein Synthesis | | | | | | Translation of protein | | | | | 0.000005 | |
| Cancer,Cell Death and Survival,Organismal Injury and Abnormalities,Tumor Morphology | | | | | | Necrosis of malignant tumor | | | | | 0.00000533 | |
| Cancer,Cell Death and Survival,Organismal Injury and Abnormalities,Tumor Morphology | | | | | | Necrosis of tumor | | | | | 0.00000799 | |
| Cell Death and Survival | | | | | | Cell survival | | | | | 0.0000118 | |
| Protein Synthesis | | | | | | Expression of protein | | | | | 0.000015 | |
| Cancer,Cell Death and Survival,Organismal Injury and Abnormalities,Tumor Morphology | | | | | | Cell death of cancer cells | | | | | 0.0000156 | |
| Cellular Development,Cellular Growth and Proliferation,Hematological System Development and Function,Lymphoid Tissue Structure and Development | | | | | | Cell proliferation of T lymphocytes | | | | | 0.0000185 | |
| Cancer,Cell Death and Survival,Organismal Injury and Abnormalities,Tumor Morphology | | | | | | Cell death of tumor cells | | | | | 0.0000421 | |
| Cellular Development,Cellular Growth and Proliferation,Connective Tissue Development and Function,Tissue Development | | | | | | Cell proliferation of fibroblasts | | | | | 0.0000504 | |
| Cell Death and Survival | | | | | | Cell viability | | | | | 0.0000554 | |
| Cell Death and Survival | | | | | | Cell viability of tumor cell lines | | | | | 0.0000594 | |
| Cellular Development,Cellular Growth and Proliferation | | | | | | Cell proliferation of tumor cell lines | | | | | 0.000219 | |
| Cell Death and Survival | | | | | | Killing of Staphylococcus aureus subsp. aureus str. Newman | | | | | 0.000254 | |
| Cellular Development,Cellular Growth and Proliferation | | | | | | Re-entry into growth of leukemia cell lines | | | | | 0.000254 | |
| Cellular Development,Cellular Growth and Proliferation,Hematological System Development and Function,Hematopoiesis,Lymphoid Tissue Structure and Development,Tissue Development | | | | | | Leukopoiesis | | | | | 0.000277 | |
| Protein Synthesis | | | | | | Metabolism of cellular protein | | | | | 0.000287 | |
| Cell-mediated Immune Response,Cellular Development,Cellular Function and Maintenance,Cellular Growth and Proliferation,Embryonic Development,Hematological System Development and Function,Hematopoiesis,Lymphoid Tissue Structure and Development,Organ Development,Organismal Development,Tissue Development | | | | | | Differentiation of T lymphocytes | | | | | 0.000306 | |
| Cellular Development,Cellular Growth and Proliferation,Hematological System Development and Function,Humoral Immune Response,Lymphoid Tissue Structure and Development | | | | | | Proliferation of B lymphocytes | | | | | 0.000325 | |
| Cell Death and Survival | | | | | | Apoptosis of tumor cell lines | | | | | 0.000327 | |
| Cellular Development,Cellular Growth and Proliferation,Embryonic Development,Hematological System Development and Function,Hematopoiesis,Lymphoid Tissue Structure and Development,Organ Development,Organismal Development,Tissue Development | | | | | | Lymphopoiesis | | | | | 0.000413 | |
| Cellular Development,Cellular Growth and Proliferation,Digestive System Development and Function,Embryonic Development,Hepatic System Development and Function,Organ Development,Organismal Development,Tissue Development | | | | | | Formation of hepatocytes | | | | | 0.000446 | |
| Cell Death and Survival | | | | | | Cell death of tumor cell lines | | | | | 0.000489 | |
| Cell-mediated Immune Response,Cellular Development,Cellular Function and Maintenance,Cellular Growth and Proliferation,Embryonic Development,Hematological System Development and Function,Hematopoiesis,Lymphoid Tissue Structure and Development,Organ Development,Organismal Development,Tissue Development | | | | | | T cell development | | | | | 0.000531 | |
| Cell Death and Survival | | | | | | Cell viability of pancreatic cancer cell lines | | | | | 0.000561 | |
| Cellular Development,Renal and Urological System Development and Function | | | | | | Differentiation of podocytes | | | | | 0.000606 | |
| Cancer,Cellular Development,Cellular Growth and Proliferation,Organismal Injury and Abnormalities,Tumor Morphology | | | | | | Proliferation of cancer cells | | | | | 0.000609 | |
| Cell Death and Survival,Organismal Injury and Abnormalities,Renal and Urological Disease | | | | | | Apoptosis of kidney cells | | | | | 0.000624 | |
| Cell Death and Survival | | | | | | Apoptosis of kidney | | | | | 0.000641 | |
| Cellular Development,Cellular Growth and Proliferation,Hematological System Development and Function,Hematopoiesis,Lymphoid Tissue Structure and Development,Tissue Development | | | | | | Proliferation of myeloblasts | | | | | 0.000755 | |
| Cell Death and Survival,Cellular Compromise | | | | | | Cytotoxicity of splenocytes | | | | | 0.000755 | |
| Cellular Assembly and Organization,Cellular Compromise,Cellular Development,Cellular Growth and Proliferation,Digestive System Development and Function,Embryonic Development,Hepatic System Development and Function,Organ Development,Organismal Development,Tissue Development | | | | | | Formation of Mallory bodies | | | | | 0.000755 | |
| Cellular Development,Cellular Growth and Proliferation,Hematological System Development and Function,Hematopoiesis,Humoral Immune Response,Lymphoid Tissue Structure and Development | | | | | | Proliferation of pro-B lymphocytes | | | | | 0.000968 | |
| Cell Death and Survival,Organismal Injury and Abnormalities,Renal and Urological Disease | | | | | | Apoptosis of tubular cells | | | | | 0.00119 | |
| Protein Synthesis | | | | | | Quantity of protein lipid complex in blood | | | | | 0.00119 | |
| Cancer,Cellular Development,Cellular Growth and Proliferation,Organismal Injury and Abnormalities,Tumor Morphology | | | | | | Proliferation of tumor cells | | | | | 0.00135 | |
| Cell Death and Survival,Organismal Injury and Abnormalities,Renal and Urological Disease | | | | | | Necrosis of renal tubule | | | | | 0.00146 | |
| Cell Death and Survival,Organismal Injury and Abnormalities | | | | | | Necrosis of epithelial tissue | | | | | 0.00221 | |
| **Top significant tox function** | | | | | | | | | | | | |
| Categories | | | Diseases.or.Functions.Annotation | | | | | | | p-value | | |
| Liver Steatosis | | | Hepatic steatosis | | | | | | | 0.000131 | | |
| Liver Steatosis | | | Microvesicular hepatic steatosis | | | | | | | 0.0077 | | |
| Liver Inflammation/Hepatitis,Liver Steatosis | | | Nonalcoholic steatohepatitis | | | | | | | 0.00886 | | |
| Liver Inflammation/Hepatitis,Liver Steatosis | | | Steatohepatitis | | | | | | | 0.0101 | | |
| Liver Steatosis | | | Nonalcoholic fatty liver disease | | | | | | | 0.0109 | | |
| Liver Steatosis | | | Methionine choline-deficient diet induced nonalcoholic fatty liver disease | | | | | | | 0.016 | | |
| Liver Steatosis | | | Severe microvesicular hepatic steatosis | | | | | | | 0.016 | | |
| Liver Steatosis | | | Advanced stage hepatic steatosis | | | | | | | 0.0472 | | |
| Liver Enlargement | | | Hepatomegaly | | | | | | | 0.00106 | | |
| Liver Proliferation | | | Proliferation of liver | | | | | | | 0.00149 | | |
| Liver Proliferation | | | Proliferation of liver cells | | | | | | | 0.00758 | | |
| Liver Fibrosis,Liver Proliferation | | | Proliferation of hepatic stellate cells | | | | | | | 0.0155 | | |
| Liver Proliferation | | | Proliferation of hepatic progenitor cells | | | | | | | 0.0317 | | |
| Liver Hyperplasia/Hyperproliferation | | | Liver tumor | | | | | | | 0.0023 | | |
| Hepatocellular carcinoma,Liver Hyperplasia/Hyperproliferation | | | Development of hepatocellular carcinoma | | | | | | | 0.00277 | | |
| Liver Hyperplasia/Hyperproliferation | | | Familial hepatic adenomas | | | | | | | 0.016 | | |
| Liver Hyperplasia/Hyperproliferation | | | Liver cancer | | | | | | | 0.0203 | | |
| Liver Hyperplasia/Hyperproliferation | | | Liver carcinoma | | | | | | | 0.0275 | | |
| Hepatocellular carcinoma,Liver Hyperplasia/Hyperproliferation | | | Unresectable hepatocellular carcinoma | | | | | | | 0.0412 | | |

**Supplemental Table 3: Epitranscriptomic analysis**

| ***Modification*** | ***tR (min)*** | ***m/z*** | ***q value*** | ***Fold Change*** | | | |
| --- | --- | --- | --- | --- | --- | --- | --- |
| **12 weeks** | | | | | | | |
|  |  |  |  | ***WD vs. CD*** | ***CD+VC vs. CD*** | ***WD+VC vs. WD*** | ***WD+VC vs. CD+VC*** |
| m2A (2-Methyladenosine) | 7.54 | 282.1199 | 0.008 | 0.83 | 1.72 | 0.83 | 0.40 |
| m2,8A | 9.73 | 296.1358 | 0.019 | 0.55 | 1.61 | 1.01 | 0.35 |
| GMP | 4.91 | 364.0657 | 0.022 | 0.79 | 0.98 | 0.83 | 0.68 |
| UMP | 3.42 | 325.0435 | 0.022 | 0.89 | 1.09 | 0.65 | 0.53 |
| m3U | 8.87 | 259.0932 | 0.022 | 0.65 | 0.90 | 1.71 | 1.22 |
| AMP | 4.70 | 348.0707 | 0.022 | 0.84 | 1.06 | 0.67 | 0.53 |
| m4Cm | 6.94 | 272.1250 | 0.027 | 1.25 | 1.24 | 0.81 | 0.82 |
| ADP | 3.75 | 428.0373 | 0.027 | 0.64 | 0.80 | 2.31 | 1.84 |
| m4C (N4-methylcytidine),m3C (3-Methylcytidine) | 5.26 | 258.1089 | 0.033 | 0.90 | 1.06 | 2.14 | 1.81 |
| m22G (N2,N2-dimethylguanosine) | 10.68 | 312.1307 | 0.038 | 0.36 | 0.42 | 1.79 | 1.56 |
| ms2m6A (2-methylthio-N6-methyladenosine) | 23.12 | 328.1080 | 0.038 | 1.86 | 1.33 | 0.77 | 1.09 |
| G (Guanosine) | 6.99 | 284.0993 | 0.038 | 0.86 | 0.76 | 1.50 | 1.69 |
| m1A (1-Methyladenosine) | 5.45 | 282.1192 | 0.038 | 0.67 | 0.96 | 1.51 | 1.06 |
| m42Cm | 7.98 | 286.1400 | 0.038 | 1.32 | 1.01 | 0.84 | 1.10 |
| mcm5s2U | 12.75 | 333.0758 | 0.038 | 0.75 | 1.02 | 1.01 | 0.74 |
| **12 months** | | | | | | | |
|  |  |  |  | ***WD vs. CD*** | ***CD+VC vs. CD*** | ***WD+VC vs. WD*** | ***WD+VC vs. CD+VC*** |
| CMP | 2.92 | 324.0595 | 0.0001 | 0.66 | 1.11 | 0.87 | 0.52 |
| m6Am | 11.33 | 296.1359 | 0.0001 | 1.49 | 1.03 | 0.94 | 1.36 |
| m2A (2-Methyladenosine) | 7.54 | 282.1199 | 0.0001 | 0.69 | 0.86 | 0.83 | 0.67 |
| m4C (N4-methylcytidine), m3C (3-Methylcytidine) | 5.26 | 258.1089 | 0.0052 | 1.79 | 0.70 | 2.95 | 7.59 |
| ADP | 3.75 | 428.0373 | 0.0116 | 1.46 | 0.76 | 1.20 | 2.32 |
| m4Cm | 6.94 | 272.1250 | 0.0116 | 2.19 | 1.00 | 0.42 | 0.92 |
| m3U | 8.87 | 259.0932 | 0.0153 | 1.07 | 0.72 | 1.17 | 1.75 |
| m8A | 8.70 | 282.1202 | 0.0153 | 0.32 | 0.88 | 0.73 | 0.26 |
| N6-SAR | 10.68 | 384.1168 | 0.0171 | 1.90 | 0.94 | 0.92 | 1.86 |
| AMP | 4.70 | 348.0707 | 0.0171 | 0.63 | 1.12 | 0.70 | 0.39 |
| UMP | 3.42 | 325.0435 | 0.0208 | 0.54 | 1.11 | 0.83 | 0.40 |
| m6A (6-Methyladenosine) | 9.04 | 282.1200 | 0.0208 | 1.44 | 1.30 | 1.46 | 1.62 |
| ac4C (N4-acetylcytidine) | 9.50 | 286.1041 | 0.0252 | 1.31 | 0.87 | 1.03 | 1.55 |
| IMP | 4.82 | 349.0547 | 0.0252 | 1.05 | 0.96 | 0.43 | 0.48 |
| **12 months vs. 12 weeks** | | | | | | | |
|  |  |  |  | ***CD 12-M/12-W*** | ***CD+VC 12-M/12-W*** | ***WD 12-M/12-W*** | ***WD+VC 12-M/12-W*** |
| m2A (2-Methyladenosine) | 7.54 | 282.1199 | 0.0000 | 1.03 | 0.52 | 0.86 | 0.85 |
| m4Cm | 6.94 | 272.1250 | 0.0001 | 1.23 | 1.00 | 2.17 | 1.12 |
| m4C (N4-methylcytidine), m3C (3-Methylcytidine) | 5.26 | 258.1089 | 0.0001 | 0.96 | 0.63 | 1.91 | 2.63 |
| Gm (2'-O-methylguanosine) | 8.75 | 298.1150 | 0.0006 | 0.63 | 0.73 | 0.74 | 0.79 |
| m2,8A | 9.73 | 296.1358 | 0.0008 | 0.61 | 0.30 | 0.90 | 0.95 |
| AMP | 4.70 | 348.0707 | 0.0008 | 0.93 | 0.99 | 0.69 | 0.72 |
| m3U | 8.87 | 259.0932 | 0.0008 | 0.83 | 0.66 | 1.39 | 0.95 |
| ADP | 3.75 | 428.0373 | 0.0008 | 0.72 | 0.69 | 1.65 | 0.86 |
| Cm (2'-O-methylcytidine) | 6.35 | 258.1091 | 0.0008 | 0.61 | 0.87 | 0.70 | 0.70 |
| m8A | 8.70 | 282.1202 | 0.0013 | 2.06 | 2.01 | 0.95 | 0.70 |
| m6A (6-Methyladenosine) | 9.04 | 282.1200 | 0.0043 | 0.65 | 1.47 | 1.21 | 1.75 |
| m2,2G (N2,N2-dimethylguanosine) | 10.68 | 312.1307 | 0.0054 | 0.40 | 1.54 | 0.90 | 0.75 |
| IMP | 4.82 | 349.0547 | 0.0078 | 1.38 | 1.27 | 1.96 | 0.85 |
| m5Um | 10.81 | 273.1089 | 0.0085 | 0.58 | 0.82 | 0.87 | 0.72 |
| m1A (1-Methyladenosine) | 5.45 | 282.1192 | 0.0086 | 0.95 | 0.79 | 1.36 | 1.40 |
| Um | 8.39 | 259.0931 | 0.0089 | 0.70 | 0.73 | 0.74 | 0.86 |
| mcm5s2U | 12.75 | 333.0758 | 0.0160 | 0.81 | 0.77 | 1.28 | 1.19 |
| m6,6A | 12.63 | 296.1360 | 0.0173 | 0.48 | 1.03 | 0.80 | 1.15 |
| Im (2'-O-methylinosine) | 8.88 | 283.1043 | 0.0173 | 0.45 | 1.75 | 0.80 | 0.82 |
| m5C (5-methylcytidine) | 5.48 | 258.1090 | 0.0208 | 0.92 | 0.56 | 1.23 | 1.07 |
| m2,7G | 7.92 | 312.1306 | 0.0246 | 1.67 | 0.86 | 1.29 | 1.19 |
| ac4C (N4-acetylcytidine) | 9.50 | 286.1041 | 0.0259 | 0.65 | 0.68 | 1.16 | 1.07 |
| ms2m6A (2-methylthio-N6-methyladenosine) | 23.12 | 328.1080 | 0.0337 | 1.90 | 1.81 | 0.80 | 1.31 |
| m2G (N2-methylguanosine) | 8.95 | 298.1148 | 0.0337 | 0.72 | 1.40 | 0.86 | 1.10 |
| DHU | 4.12 | 247.0928 | 0.0433 | 0.95 | 0.87 | 1.70 | 0.99 |

C. **Figures:**

**Supplemental Figure 1:** Immunohistochemistry was performed for Ki-67 (cellular proliferation) and CD31 (angiogenesis). VC significantly increased Ki-67 positive staining independent of diet. No changes were observed for CD31.

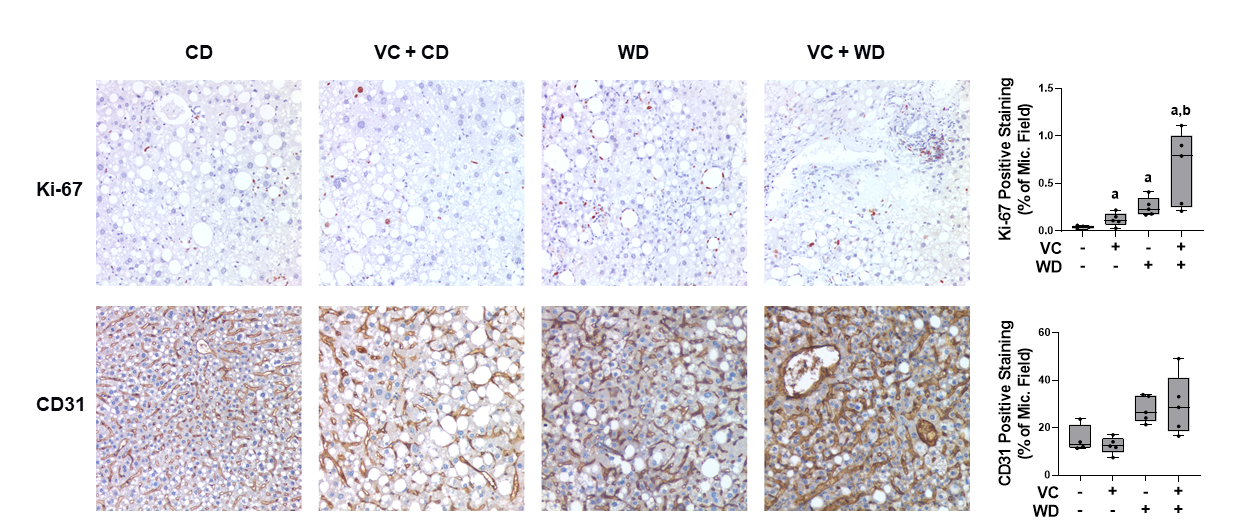

**Supplemental Figure 2:** Liver sections were stained, or immunohistochemistry was performed for Oil Red O (ORO, lipids), 4-hydroxynonenal (4-HNE, oxidative stress) and Sirius Red (SR, collagen). Cholesterol and cholesterol esters were measured in hepatic lipid extracts. While WD increased all of these indices, VC did not increase these effects, except for SR. VC increased SR positive staining in the WD group.

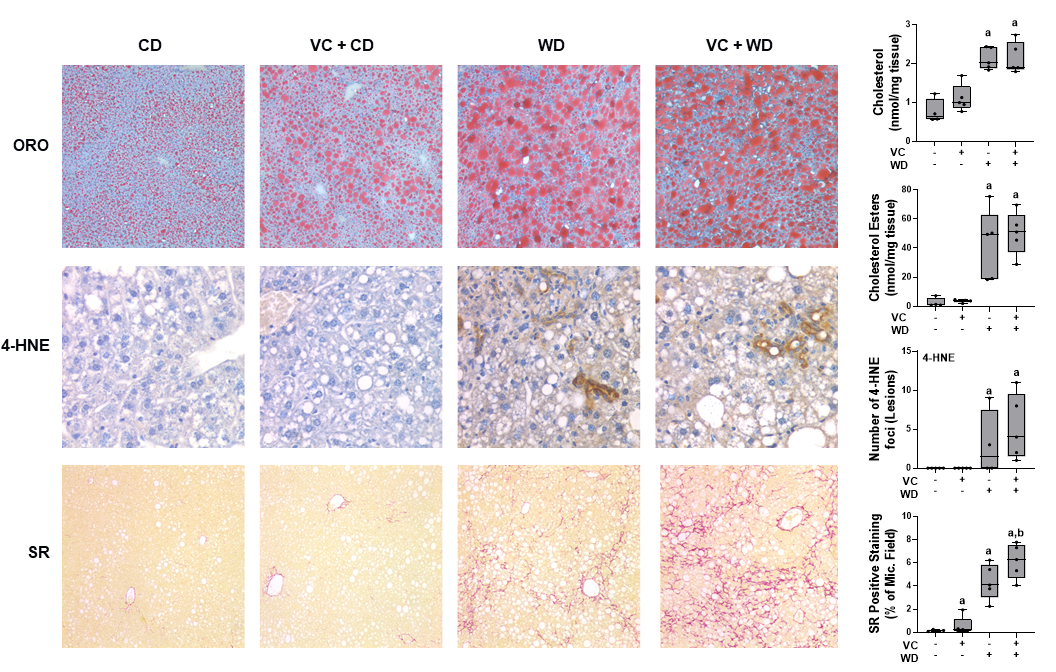

**Supplemental Figure 3:** Left panel: top KEGG, GO, and GO BP terms identified changed by IPA analysis Right panel: volcano plot comparing the expression pattern in mice exposed to VC (versus air) at the 12-week timepoint. The red dots represent significantly up- or down-regulated genes, and the grey dots represent insignificant differentially expressed genes.

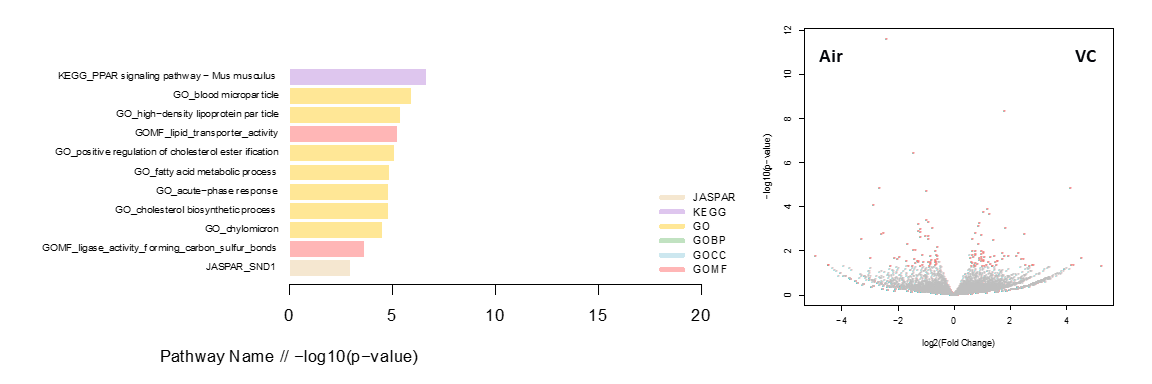

**Supplemental Figure 4:** Top pathways identified changed by IPA analysis.

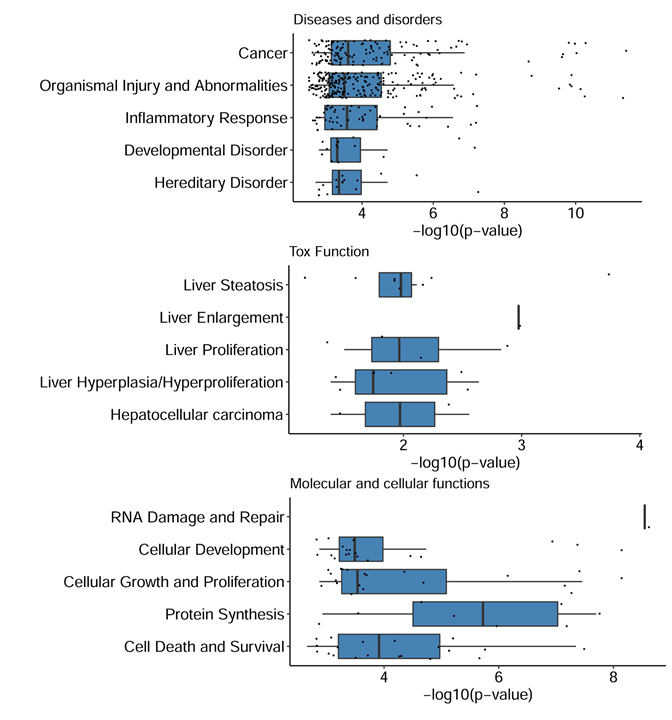

**Supplemental Figure 5:** PCA and PLS-DA analysis of total hepatic nucleosides and nucleotides at 12 weeks

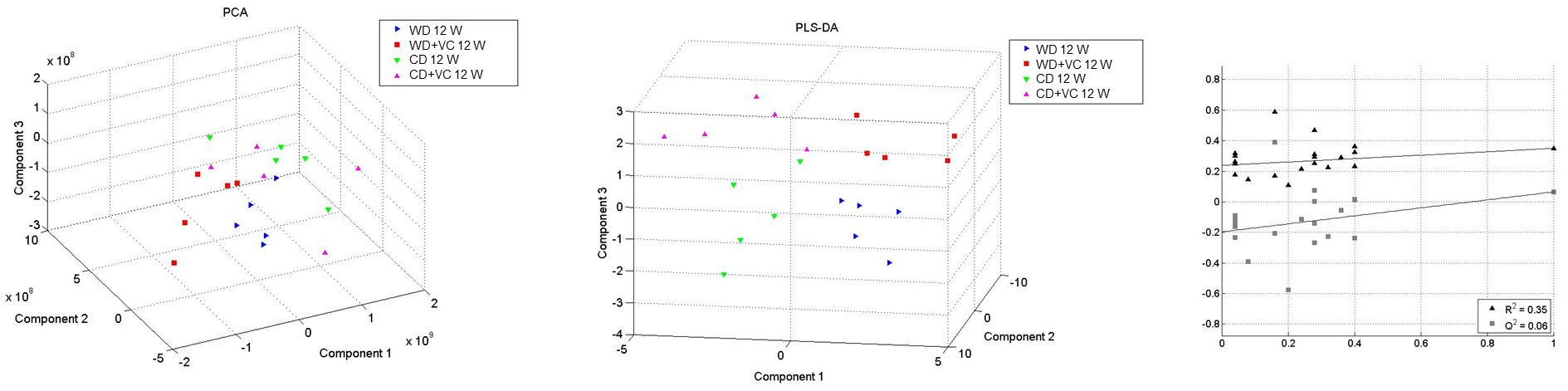

**Supplemental Figure 6:** LC-MS/MS epitranscriptomic analysis of total hepatic nucleosides and nucleotides at 12 weeks VC vs. air. *, p<0.05 log2 fold change (FC). n=5 per group.

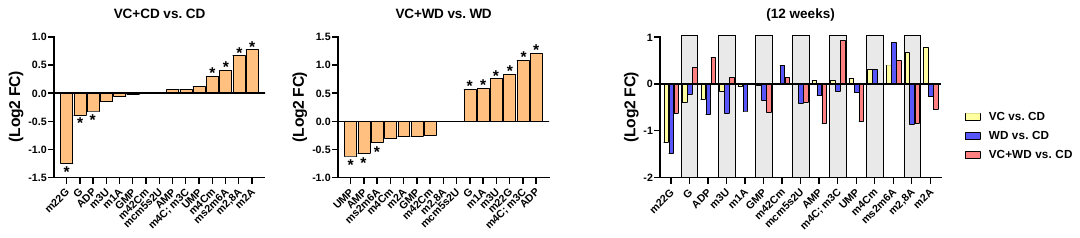

**Supplemental Figure 7:** PCA and PLS-DA analysis of total hepatic nucleosides and nucleotides at 12 months

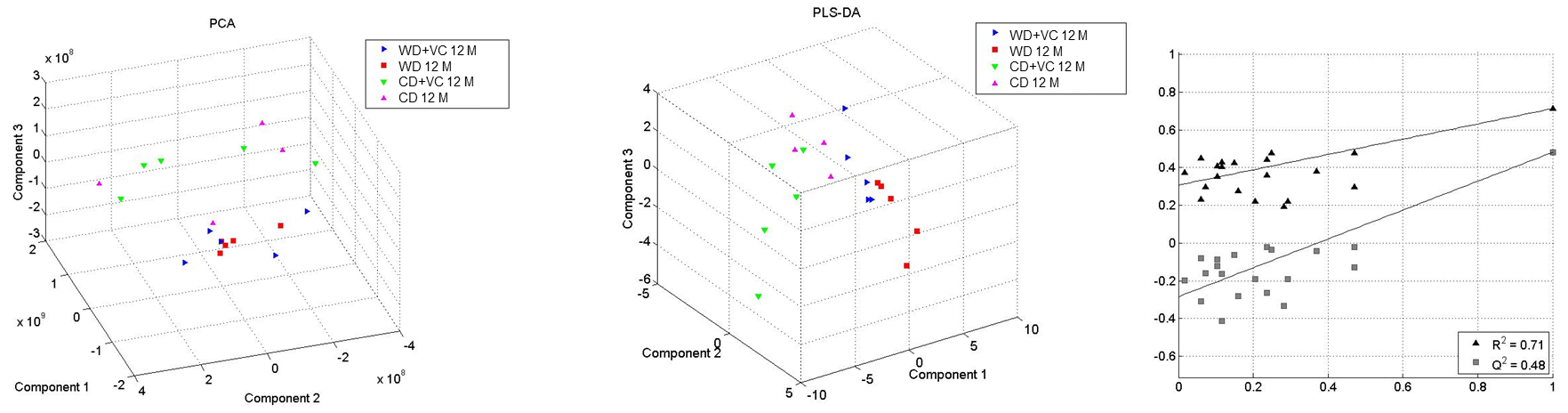

**Supplemental Figure 8:** PCA and PLS-DA analysis of total hepatic nucleosides and nucleotides 12 months vs 12 weeks

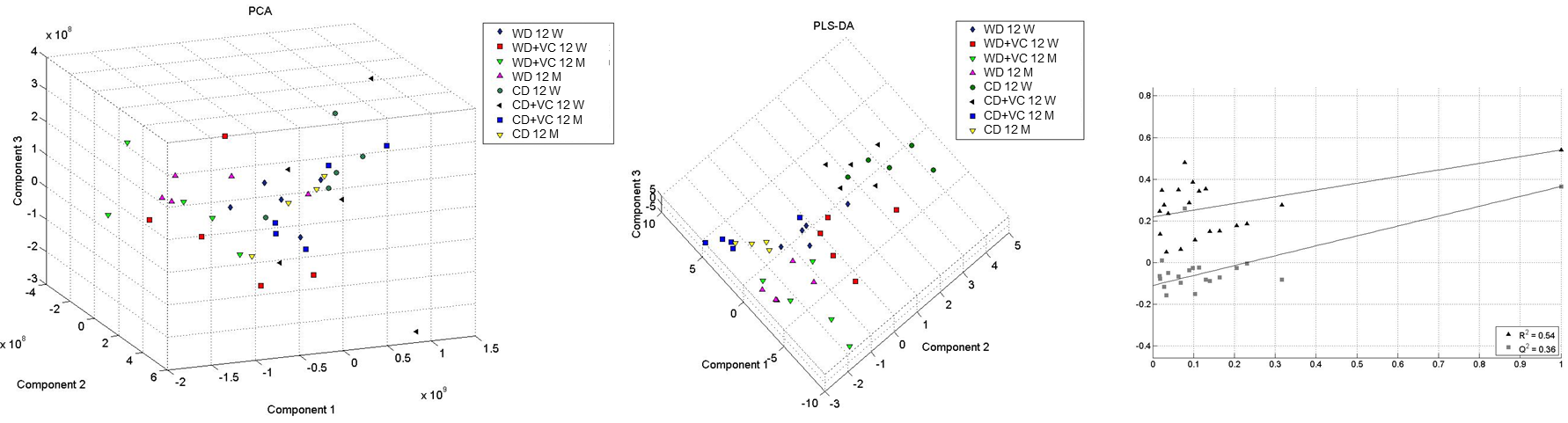
